# Insights into the genetic architecture of complex traits in Napier grass (*Cenchrus purpureus*) and QTL regions governing forage biomass yield, water use efficiency and feed quality traits

**DOI:** 10.1101/2021.03.15.435454

**Authors:** Meki S. Muktar, Ermias Habte, Abel Teshome, Yilikal Assefa, Alemayehu T. Negawo, Ki-Won Lee, Jiyu Zhang, Chris S. Jones

## Abstract

Napier grass is the most important perennial tropical grass native to Sub-Saharan Africa and widely grown in tropical and subtropical regions around the world, primarily as a forage crop for animal feed, but with potential as an energy crop and in a wide range of other areas. Genomic resources have recently been developed for Napier grass that need to be deployed for genetic improvement and molecular dissection of important agro-morphological and feed quality traits. From a diverse set of Napier grass genotypes assembled from two independent collections, a subset of 84 genotypes that best represents the genetic diversity of the collections were selected and evaluated for two years in dry (DS) and wet (WS) seasons under three soil moisture conditions: moderate water stress in DS (DS-MWS); severe water stress in DS (DS-SWS) and, under rainfed (RF) conditions in WS (WS-RF). Data for agro-morphological and feed quality traits, adjusted for the spatial heterogeneity in the experimental blocks, were collected over a two-year period from 2018 to 2020. A total of 135,706 molecular markers were filtered, after removing markers with missing values > 10% and a minor allele frequency (MAF) < 5 %, from the high-density genome-wide markers generated previously using the genotyping by sequencing (GBS) method of the DArTseq platform. A genome-wide association study (GWAS), using two different mixed linear model algorithms, identified more than 58 QTL regions and markers associated with agronomic, morphological and water-use efficiency traits. QTL regions governing purple pigmentation and feed quality traits were also identified. The identified markers will be useful in the genetic improvement of Napier grass through the application of marker-assisted selection and for further characterization and map-based cloning of the QTLs.

## 1 Introduction

Improving livestock feeds and forages will play a key role in global food and nutrition security and have the potential to contribute to the strategy of achieving climate-smart agriculture, restoring degraded lands and decreasing greenhouse gas emission intensities (Paul et al., 2020; Belay 2019; Bryan et al., 2013; Peters et al., 2013). The availability of adequate, high-quality feeds and forages has been a major challenge faced by the livestock sector, especially during the dry season when pasture and crop residues are scarce (Maleko et al., 2018; Mtengeti et al., 2008). To cope with the shortage of feeds during the dry season, many farmers in sub-Saharan Africa (SSA) rely mainly on drought-tolerant perennial grasses, such as Napier grass, that can produce a reasonable amount of feed under limited water availability (Kabirizi et al., 2015; Lukuyu et al., 2012).

Napier grass (*Cenchrus purpureus* (Schumach.) Morrone syn. *Pennisetum purpureum* Schumach.), also called elephant grass, belonging to the poaceae family, is one of the most important perennial tropical C4 grasses (Robert et al., 2010). Napier grass is native to SSA from where it has been distributed to other tropical and subtropical regions around the world, adapting to a wide range of soil and agro-ecological conditions (Negawo et al., 2017). Napier grass has been adapted to areas of North and South America, tropical parts of Asia, Australia, the Middle East, and, the Pacific (Negawo et al., 2017; Anderson et al., 2008). Napier grass is cultivated primarily as a forage crop for animal feed in cut-and-carry feeding systems, it is particularly well-known by smallholder farmers in Eastern and Central Africa (Kabirizi et al., 2015; Lukuyu et al., 2012). Napier grass is known for its high biomass production (up to 78 tons of dry matter per hectare annually), year-round availability under limited irrigation, ability to withstand repeated cuttings when harvested multiple times, resistance to most pests and diseases, ease of establishment and rapid propagation and, fast regrowth capacity (Kabirizi et al., 2015; Lukuyu et al., 2012; Anderson et al., 2008). Napier grass is also used in the push-pull integrated pest management strategy (Van den Berg et al., 2015; Khan et al., 2011), is commonly grown around many crops as a wind and fire break and, is planted in marginal lands and slopes to increase soil fertility and to reduce soil erosion (Negawo et al., 2017; Kabirizi et al., 2015). Recently, reports have shown the potential of Napier grass for biofuel, bioremediation and paper production (Rocha-Meneses et al., 2020; Tsai and Tsai, 2016; Rengsirikul et al., 2013; Madakadze et al., 2010).

Napier grass is an allotetraploid (2n = 4x = 28) with a complex genome (A'A'BB genomes, with A’ genome showing a high degree of homology with the pearl millet A genome) and high genetic diversity, which is mainly attributed to the wide parental diversity and its out-crossing nature (Robert et al., 2010; dos Reis et al., 2014). Despite its multipurpose use, enormous potential in a wide range of areas and its high genetic diversity, there have been limited efforts to develop varieties with high forage value through breeding and genetic studies. Most of the genetic studies conducted so far were limited to characterizing its genetic diversity using low-density molecular markers (Negawo et al., 2018; Kandel et al., 2016; Babu et al., 2009; Wanjala et al., 2013; Azevedo et al., 2012) that provide a poor representation of the whole-genome information. The first detailed genetic diversity and population structure using genome-wide representative markers was the report by Muktar et al. (2019), in which high-density genome-wide single nucleotide polymorphism (SNP) and SilicoDArT markers were generated using the genotyping by sequencing (GBS) method of the DArTseq platform. This study provided a useful insight into the genetic diversity and genome-wide patterns of linkage disequilibrium (LD) in the Napier grass collections, both the material maintained in the International Livestock Research Institute (ILRI) forage genebank and a collection acquired from the Brazilian Agricultural Research Corporation (EMBRAPA), and demonstrated the potential of the collections for further genetic and marker-trait association studies. The generation of high-density genome-wide markers also permitted the construction of the first high-density genetic map of Napier grass (Paudel et al., 2018). The most important breakthrough was the recent reports of the first high-quality chromosome scale genome sequences of Napier grass (Yan et al., 2021; Zhang et al., 2020), in which a 1.97 to 2.07 Gb genome was assembled. This chromosome scale genome sequence offers significant opportunities for the dissection of the genetic architectures of complex traits and the development of improved Napier grass varieties.

Molecular tools need to be deployed for Napier grass improvement and the dissection of important agronomic and feed quality traits, for example, by linking sequence polymorphisms with traits using genome-wide association studies (GWAS) and/or linkage mapping. The GWAS technique, which is based on LD, is well established and a potential approach for genetic dissection of complex traits and the identification of QTLs (Huang and Han, 2014; Flint-Garcia et al., 2003). DNA markers identified by GWAS for agronomic traits have been successfully exploited in several crop plants, for marker-assisted selection (MAS), gene cloning, trait improvement and designing an effective breeding strategy (Jaiswal et al., 2019; Guo et al., 2018; Sanchez et al., 2018; Burridge et al., 2017). Unlike classical linkage mapping that uses a population of the progeny of a biparental cross, GWAS is performed on a diverse collection of unrelated genotypes (Huang and Han, 2014; Flint-Garcia et al., 2003) and hence this technique is ideally suited to the study of genebank collections (McCouch et al., 2012). To date, only two GWAS studies on Napier grass have been reported (Habte et al., 2020; Rocha et al., 2019), in which markers associated with high biomass yield, metabolizable energy and biomass digestibility were detected. However, these studies had a limitation on either marker density or population size.

Here, we report on the first marker-trait association and QTL identification in Napier grass in a controlled and replicated field trial, with repeated trait measurements, high-density genome-wide markers and thorough statistical approaches. Selected Napier grass genotypes have been evaluated in a field in dry (DS) and wet (WS) season conditions. Morphological, agronomic, water-use efficiency and feed quality traits were collected over two years from 2018 to 2020, the trait values were subjected to a spatial analysis (Rodríguez-Álvarez et al., 2018) and adjusted for the spatial heterogeneity of the experimental blocks. GWAS was employed using the adjusted phenotypic data and high-density genome-wide markers generated previously (Muktar et al., 2019), with the objective of dissecting the genetic architecture of complex traits in Napier grass and identifying markers and QTL regions associated with forage-biomass yield, water-use efficiency and feed quality traits.

## 2 Materials and Methods

### Selection of high-density genome-wide markers

The Napier grass collections have been genotyped as described previously (Muktar et al., 2019). From the high-density genome-wide markers generated, a total of 135,706 markers (90,498 silicoDArTs and 45,208 SNPs) were filtered after removing markers with missing values > 10% and a minor allele frequency (MAF) < 5 %. The markers with 10 % missing values were imputed using the missForest r package (Stekhoven and Bühlmann, 2012), with maxiter set to 5 and ntree to 100 and all other parameters set to default values. The imputation was run on one assembled chromosome (AC) at a time as the run time of the software could not accommodate taking the entire genome at once. The short sequence reads corresponding to the markers were aligned to the recently reported Napier grass genome (Yan et al., 2021) using the bwa mem sequence aligner (Li and Durbin, 2009) and the marker density and distribution across the fourteen ACs of the Napier grass genome was visualized using the synbreed R-package (Wimmer et al., 2012). The sequences were annotated using the genomic information resources of *Cenchrus americanus* and *Setaria italica* in the GenBank NCBI blastx tool (https://blast.ncbi.nlm.nih.gov/Blast.cgi) as described previously (Muktar et al., 2019).

### Marker data analysis, linkage disequilibrium (LD) and LD-decay analysis

The missing percentage data, minor allele frequency (MAF) per marker and per genotype, and the polymorphic information content (PIC) were calculated in R statistical software (https://www.r-project.org/) as described previously (Muktar et al., 2019). Pair-wise linkage disequilibrium (LD) between pairs of SilicoDArT markers (SilicoDArTs were selected as the DArTseq technology produces more precise genomic position information for this marker than for SNPs) with a known genomic location on the Napier grass genome (Yan et al., 2021) was analysed. The marker data analysis, LD and LD-decay across the Napier grass genome and population structure analysis can be found in Muktar et al. (2019).

### Field planting, drought stress application and trait measurements

A total of 84 genetically diverse genotypes were evaluated at the Bishoftu field site, Ethiopia. The genotypes were arranged in a partially replicated (p-rep) design in four blocks as described previously (Muktar et al., 2019). Six plants per accession were planted in a single row, with 750 mm spacing between plants and between rows. During the dry season (DS), two blocks were irrigated to a volumetric soil water content (VWC) of approximately 20% (now onwards called moderate water stress, MWS) and the other two blocks were irrigated with a reduced amount of water, which corresponds to a VWC of about 10% (now onwards called severe water stress, SWS) (Supplementary Figure 1). There was no irrigation in the wet season (WS) as all plants were maintained under rainfed (RF) conditions (approximately 30% VWC) (Supplementary Figure 1 and 2). Overall, 12 harvests were conducted, following every eight weeks of regrowth, in both WS and DS conditions. Data of morphological, agronomic, and feed quality traits were collected from three randomly selected plants per row in each of the three soil moisture conditions, as described in (Habte et al., 2020). The agro-morphological traits collected were plant height (PH) in centimeter (cm), leaf length (LL) in cm, leaf width (LW) in millimeter (mm), stem thickness (ST) in mm, tiller number (TN) count, internode length (IL) in cm, total fresh weight (TFW) in t/ha, total dry weight (TDW) t/ha, leaf-stem-ratio (LSR), and water use efficiency (WUE) in g/l (Dry matter produced per liter of water). Acid detergent fibre (ADF) in %, acid detergent lignin (ADL) in %, crude protein (CP) in %, dry matter (DM), in vitro organic matter digestibility (IVOMD), metabolizable energy (Me) in J/KgDM, neutral detergent fibre (NDF) in %, organic matter (OM) in %, were collected for the feed quality traits. Additional information on materials and methods is found in Supplementary text 1.

### Phenotypic data analysis and correction for spatial variation

The phenotypic value of each trait was adjusted according to the spatial variation across the experimental field using the SpATS r-package (Rodríguez-Álvarez et al., 2018) in R (R Development Core Team 2016) statistical software. The analysis was performed individually for each of the three soil moisture conditions. In this study, plots were laid out in 24 row by 4 column grids in each block, thus rows and columns were used as random factors. In addition, soil moisture data and soil nutrient parameters (Acid = soil acidity, AvaP = available phosphorus, K = available potassium, OM = organic matter and CEC = cation exchange capacity) were included in the mixed model as fixed covariates.

The pairwise correlation between all possible trait-pairs was assessed using the R function “*cor_pmat*” in the package ggcorrplot in R (R Development Core Team 2020) and visualization of the correlation matrices was undertaken using the ‘*ggcorrplot*’ function. Effective dimension (ED), which is a measure of the complexity of the model components, and broad-sense heritability based on the adjusted data were generated by the r functions “*summary*” and “*getHeritability*”, respectively, in the SpATS r-package (Rodríguez-Álvarez et al., 2018). For comparison purposes, broad-sense heritabilities were also analysed for the unadjusted data using the “*mmer*” function in the sommer r package (Covarrubias-Pazaran, 2016). The normality of the data for each trait was tested by drawing normal plots in a histogram by using the “*hist*” function in R (R Development Core Team 2020).

### Marker trait association analysis

Marker-trait association analysis was carried out using 83 Napier grass genotypes (one genotype was excluded because of high missing values for the marker data) that had been genotyped and phenotyped (adjusted for spatial variation as described above).

For the agro-morphological and feed quality quantitative traits, the analysis was performed using the recently developed method Bayesian-information and Linkage-disequilibrium Iteratively Nested Keyway in R (BLINK.R) (Huang et al., 2018) and the mixed linear model (MLM) in the R package GWASpoly (Rosyara, et al., 2016), which takes population structure and kinship into account in a mixed model to correct against spurious associations due to the level of relatedness between genotypes. The BLINK model is based on linkage disequilibrium (LD) and eliminates confounding issues arising due to population structure, kinship, multiple testing correction, etc. The first three to five components identified through principal component analysis (PCA) using the Adegenet r package (Jombart, 2008) were included as covariates in the model. The number of components increased from three to five until the quantile-quantile (Q-Q) plot shows a similar distribution between observed and expected *P*-values along a solid diagonal line except for a sharp curve of the observed *P*-value at the end of the line, which represents a true association. The MLM model was based on a Q+K MLM with biallelic markers that represent population structure and relatedness, as implemented in the R package GWASpoly (Rosyara et al., 2016). From the eight different gene action models provided by the model, the additive model with ploidy = 2 was used in this study. The first five principal components and the kinship matrix, both calculated with the GWASpoly built-in algorithms, were included as covariates. To control for type I errors due to multiple testing, the *p*-values were adjusted following a false discovery rate (FDR) correction procedure (Benjamini and Hochberg, 1995). A marker with a *P*-value of less than 1.00E-05 in the BLINK model and an FDR corrected *P*-value of 0.01 in the MLM was claimed to be associated. Markers detected only by the MLM at a *P*-value of less than 1.00E-04 were selected as additional information in the supplementary table.

In addition, marker-trait association analysis was carried out for the qualitative trait, purple pigmentation, by computing the non-parametric univariate Fisher's exact test (Warner, 2013) using the Adegenet r package (Jombart, 2008). Out of the diverse set of Napier grass genotypes acquired from EMBRAPA (Muktar et al., 2019; Negawo et al., 2018), seven had purple-coloured leaves, midribs, petioles and stems (Supplementary Figure 3). The plant colours were qualitatively scored with a “1” for the seven purple-coloured genotypes and “0” for 98 green-coloured genotypes. A threshold level at *P-value* < 1.00E-07 (> the −log10 of 7) was used to claim an association.

The genomic map position of associated markers and their co-localization was estimated based on the sequence length of each of the 14-assembled chromosomes (AC) (Yan et al., 2021) and a physical map was constructed at a 1Mbp scale.

## 3 Results

### Spatially corrected phenotype data, distribution, and correlation of traits

The phenotypic measurements of the agronomic, morphological, and feed quality traits were collected on a genetically diverse set of 84 Napier grass genotypes evaluated over a two-year period under three soil moisture conditions in the wet (WS) and dry (DS) seasons. All plants in the four blocks were maintained under rainfed (RF) condition during the WS, while in the DS, two blocks were maintained under moderate water stress (MWS) and the other two blocks were maintained under severe water stress (SWS) condition using drip irrigation. The average soil volumetric water content (VWC) of each of six harvests under the three soil moisture conditions (WS-RF, DS-MWS and DS-SWS) is shown in Supplementary Figure 1. The phenotypic values were corrected for spatial heterogeneity across rows and columns of the experimental blocks, as well as for the heterogeneity in soil moisture content (SM) and soil nutrient parameters. The depicted spatial trend representing the estimated heterogeneity across rows and columns of the experimental blocks for each treatment is shown in Figure 1.

**Figure 1.**
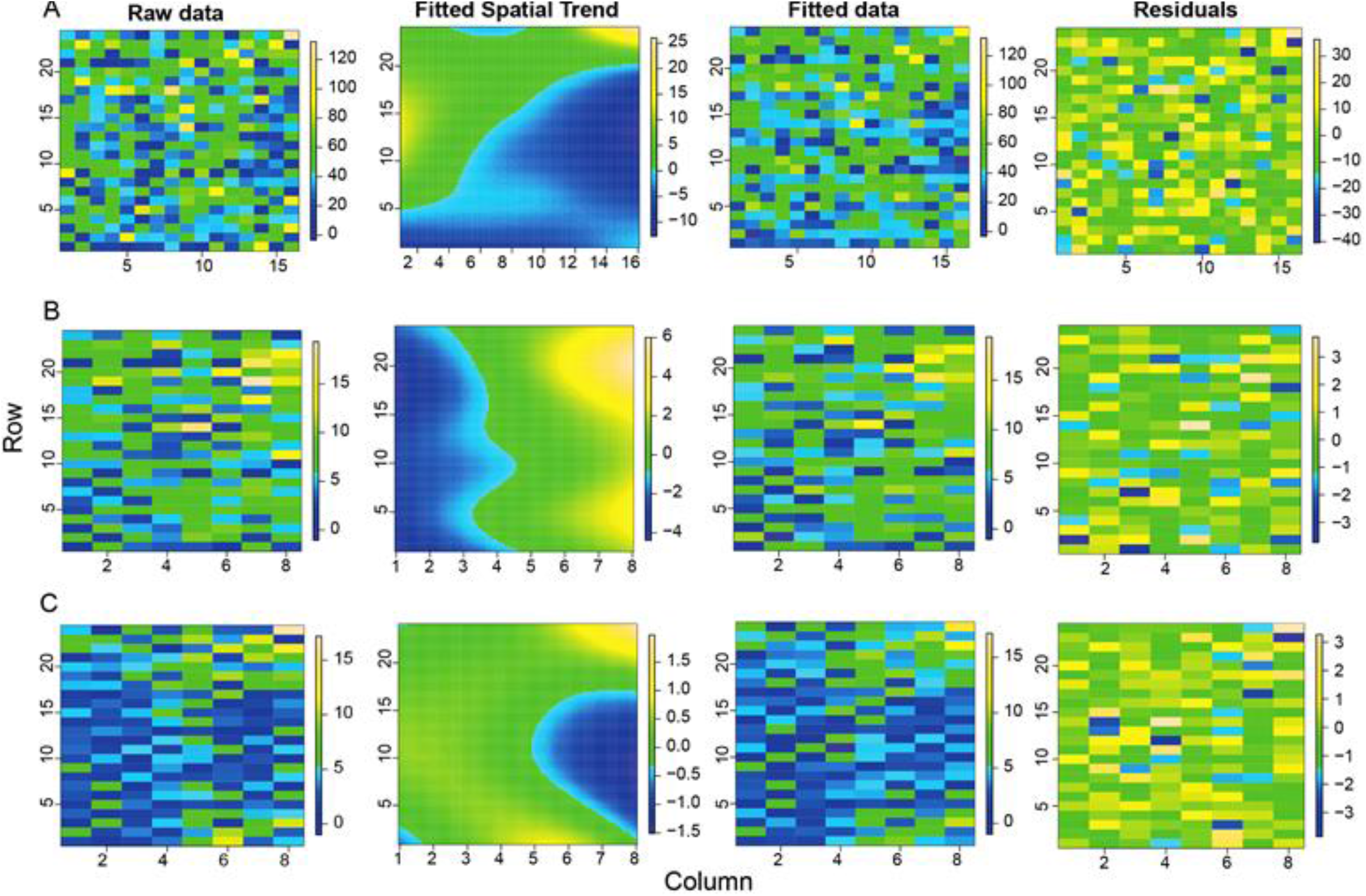
Graphical representations of the raw data, fitted spatial trend, fitted data and residuals for the biomass trait (Total fresh weight, TFW) of the wet season under rainfed (WS-RF) (A), under moderate water stress in dry season (DS-MWS) (B), and severe water stress in dry season (DS-SWS) (C) conditions.

The model captured more spatial heterogeneity towards the columns than the rows as suggested by effective dimension (ED) (Table 1), which is a measure of the complexity of the SpATS model (Rodríguez-Álvarez et al., 2018). This column-wise spatial heterogeneity was more pronounced in the WS (four blocks) than in the DS (two blocks in each treatment). As shown by the spatial trend (Figure 1A and C), plants grown in the middle part of block 4 performed worse compared to the plants in the other blocks which was consistent with visual observations. In the DS, of the two blocks under MWS conditions, plants in block 3 performed better than in block 1 (Figure 1B).

**Table 1.** Estimated effective dimension (ED) associated with the spatial trend of the row (R), column (C) and genetic (G) random factors for each trait. Broad-sense heritability for unfitted (H^2^g) and fitted (H^2^gf) data for the wet season under rainfed (WS-RF), dry-season under moderate water stress (DS-MWS) and dry season under severe water stress (DS-SWS) conditions are shown.

| Trait | WS-RF |  |  |  |  | DS-MWS |  |  |  |  | DS-SWS |  |  |  |  |
| --- | --- | --- | --- | --- | --- | --- | --- | --- | --- | --- | --- | --- | --- | --- | --- |
| | ED (C) | ED (R) | ED (G) | $H^2_g$ | $H^2_{gf}$ | ED (C) | ED (R) | ED (G) | $H^2_g$ | $H^2_{gf}$ | ED (C) | ED (R) | ED (G) | $H^2_g$ | $H^2_{gf}$ |
| PH | 10.5 | 0.0 | 73.9 | 0.80 | 0.89 | 0.8 | 1.3 | 71.1 | 0.59 | 0.86 | 0.0 | 0.0 | 61.5 | 0.31 | 0.74 |
| LL | 10.1 | 0.0 | 74.9 | 0.85 | 0.90 | NA | NA | NA | NA | NA | 2.6 | 0.0 | 57.1 | 0.41 | 0.69 |
| LW | 8.7 | 5.6 | 75.5 | 0.82 | 0.91 | 1.1 | 2.3 | 73.6 | 0.60 | 0.89 | 0.0 | 0.0 | 61.1 | 0.36 | 0.74 |
| ST | 4.3 | 0.0 | 24.7 | 0.19 | 0.30 | 1.9 | 0.0 | 46.9 | 0.49 | 0.57 | 2.5 | 0.0 | 45.6 | 0.40 | 0.56 |
| TN | 0.0 | 6.9 | 71.7 | 0.79 | 0.86 | 0.0 | 8.4 | 71.1 | 0.89 | 0.86 | 0.0 | 0.8 | 67.7 | 0.81 | 0.82 |
| IL | 8.1 | 0.1 | 51.5 | 0.29 | 0.62 | NA | NA | NA | NA | NA | NA | NA | NA | NA | NA |
| LSR | 0.2 | 0.1 | 10.4 | 0.06 | 0.12 | NA | NA | NA | NA | NA | NA | NA | NA | NA | NA |
| TFW | 5.0 | 0.1 | 69.2 | 0.78 | 0.83 | 1.8 | 0.2 | 68.2 | 0.78 | 0.82 | 1.0 | 0.0 | 58.7 | 0.66 | 0.71 |
| TDW | 0.1 | 0.0 | 58.4 | 0.51 | 0.70 | 2.5 | 0.0 | 66.2 | 0.72 | 0.80 | 0.4 | 0.0 | 58.9 | 0.69 | 0.71 |
| WUE | NA | NA | NA | NA | NA | 2.7 | 0.0 | 65.2 | 0.63 | 0.79 | 2.5 | 0.0 | 58.6 | 0.63 | 0.71 |
| ADF | 11.3 | 0.0 | 66.1 | 0.52 | 0.70 | 0.0 | 0.9 | 59.4 | 0.49 | 0.62 | 0.0 | 8.1 | 51.0 | 0.38 | 0.51 |
| ADL | 9.1 | 0.0 | 60.1 | 0.40 | 0.62 | 0.0 | 0.1 | 72.1 | 0.66 | 0.77 | 0.0 | 6.2 | 59.8 | 0.49 | 0.62 |
| CP | 7.4 | 1.0 | 44.0 | 0.23 | 0.43 | 2.1 | 0.0 | 61.0 | 0.59 | 0.64 | 0.0 | 3.9 | 53.7 | 0.29 | 0.55 |
| ash | 7.0 | 0.1 | 69.4 | 0.63 | 0.74 | 2.7 | 3.9 | 68.6 | 0.73 | 0.73 | 4.0 | 6.4 | 48.9 | 0.53 | 0.59 |
| DM | 8.2 | 0.0 | 48.8 | 0.28 | 0.49 | 2.0 | 2.1 | 70.9 | 0.70 | 0.75 | 0.0 | 0.0 | 62.1 | 0.48 | 0.65 |
| IVOMD | 8.2 | 0.0 | 48.5 | 0.27 | 0.48 | 1.1 | 0.0 | 58.3 | 0.52 | 0.60 | 0.0 | 2.0 | 47.9 | 0.23 | 0.48 |
| Me | 10.9 | 0.0 | 38.9 | 0.22 | 0.37 | 0.3 | 0.6 | 53.1 | 0.51 | 0.47 | 0.1 | 6.0 | 32.5 | 0.24 | 0.39 |
| NDF | 4.5 | 0.0 | 62.2 | 0.40 | 0.65 | 0.0 | 0.1 | 69.5 | 0.70 | 0.74 | 2.7 | 0.0 | 63.7 | 0.59 | 0.67 |
| OM | 6.5 | 0.0 | 67.2 | 0.60 | 0.71 | 2.7 | 3.9 | 68.6 | 0.73 | 0.73 | 4.0 | 0.0 | 52.6 | 0.50 | 0.53 |

PH=plant height, LL=leaf length, LW=leaf width, ST=stem thickness, TN=tiller number, IL=internode length, LSR=leaf stem ratio, TFW= total fresh weight, TDW= total dry weight, WUE=water-use efficiency, ADF=acid detergent fibre, ADL=acid detergent lignin, CP=crude protein, DM=dry matter, IVOMD=in vitro organic matter digestibility, Me=metabolizable energy, NDF=neutral detergent fibre, OM=organic matter.

Correcting the phenotypic values according to the spatial heterogeneity improved the precision of the heritability estimates and mostly increased the heritability value of the traits (Table 1), indicating the importance of the spatial analysis to determine the genetic and environmental effects on the phenotypic response and to reduce the environmental effects and errors. The heritability of traits ranged from 0.12 to 0.91 in the WS, 0.57 to 0.89 in the DS under MWS and 0.39 to 0.82 in the DS under SWS conditions. Traits LW, LL, PH, TN, ash, TFW, OM and ADF had the highest heritability followed by NDF, ADL, TDW and WUE, while ST, Me and IL had low to medium heritabilities. LSR had the lowest heritability, which indicates that the variation in the trait was mainly due to environmental factors. Therefore, the LSR trait was excluded from the GWAS analysis. With LSR excluded, heritability generally decreased in the DS, particularly under the SWS condition (Table 1).

In the correlation analysis, TFW, TDW and WUE showed a strong positive correlation (> 0.97), while LSR was strongly negatively correlated with all the traits except with IL. TN had a weak negative correlation with IL and LW, while IL showed a weak to moderate negative correlation with LW, TN, PH, TFW and TDW (Supplementary Figure 4). The phenotypic values of all the traits followed a normal distribution, only the distribution of LL under the WS-RF and DS-SWS conditions and ST and LW in the WS-RF were slightly skewed to the left (Supplementary Figure 5).

From the feed quality traits, IVOMD and Me and IVOMD and CP showed a strong positive correlation (> 0.90) followed by a strong correlation between Me and CP, ADL and ADF and NDF and OM. Conversely, OM and ash showed a very strong negative correlation (> −0.97), followed by a strong negative correlation between IVOMD and ADF, Me and ADF, NDF and ash, and, ADF and CP (Supplementary Figure 6). The values of all the traits showed a normal distribution (Supplementary Figure 7).

There were no strong positive/negative correlations between traits from the agro-morphological and feed quality traits. However, most of the agro-morphological traits, except TN, were reasonably positively correlated (0.30 to 0.60) with the traits linked with fibre components (ADF and NDF) and negatively with traits positively affecting the feed nutritional quality (CP, IVOMD and Me).

### Genome-wide distribution and density of markers on the Napier grass genome

Out of more than 200,000 high density genome-wide SilicoDArT and SNP markers (Muktar et al., 2019) generated on the Napier grass collections, a total of 135,706 (90,498 silicoDArTs and 45,208 SNPs) markers were mapped on to the Napier grass genome (Yan et al., 2021) and their distribution across the fourteen assembled chromosomes (AC) is shown in Figure 2. Approximately 80 % of the markers were mapped on the genome. The highest number of markers mapped onto A01 and B01, while the lowest number mapped onto B07 (Figure 2), with a strong correlation (> 0.94) between number of markers and chromosome size (Table 2). Polymorphic information content (PIC) values of the markers ranged from 0.08 to 0.38 with an average of 0.27 and more than 63 % of the markers had a PIC value above 0.25.

**Figure 2.**
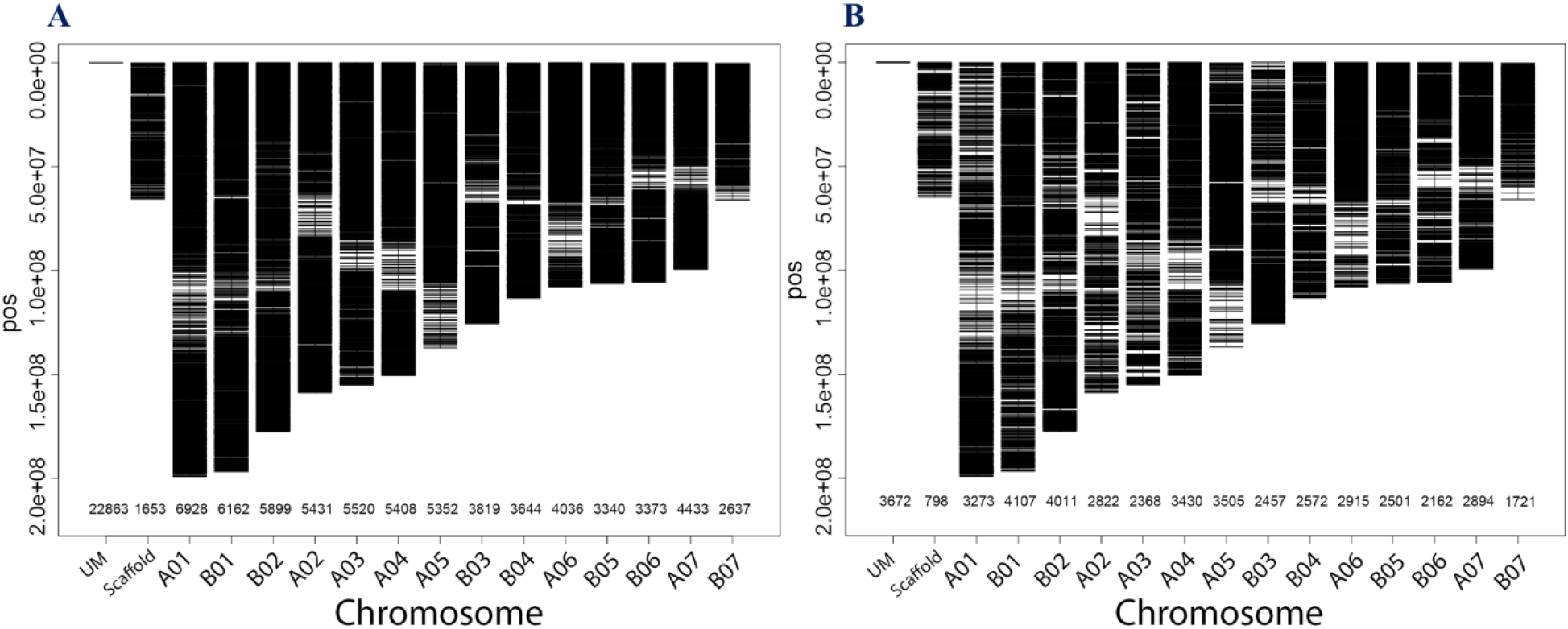
Distribution and density of SilicoDArTs (A) and SNPs (B) on the fourteen assembled chromosomes (AC) of the Napier grass genome. The number of markers mapped per AC is shown on the x-axis. The markers that were not mapped are indicated by a UM and those markers that were mapped onto different scaffolds are indicated by Scaffold.

**Table 2.** Density and distribution of markers, proportion of pairwise marker in linkage disequilibrium (LD) and the average LD-decay distance in each of the 14 assembled chromosomes (AC).

| AC | AC-size<sup>i</sup><br>(Mbp) | Number of<br>markers | markers<br>density<br>(per Mbp) | Proportion of<br>marker-pairs in<br>LD ($r^2 \geq 0.2$) | Proportion of<br>marker-pairs in<br>LD ($r^2 \geq 0.7$)<sup>ii</sup> | LD-decay<br>distance<br>(kbp) |
| --- | --- | --- | --- | --- | --- | --- |
| A01 | 199.06 | 6928 | 34.80 | 15.98 | 2.58 | 7.58 |
| B01 | 196.76 | 6162 | 31.32 | 13.61 | 1.78 | 4.61 |
| B02 | 177.74 | 5899 | 33.19 | 13.01 | 1.58 | 2.68 |
| A02 | 158.81 | 5431 | 34.20 | 14.45 | 2.37 | 5.47 |
| A03 | 155.16 | 5520 | 35.58 | 14.38 | 2.37 | 5.79 |
| A04 | 150.59 | 5408 | 35.91 | 13.43 | 1.73 | 3.85 |
| A05 | 137.44 | 5352 | 38.94 | 13.19 | 1.78 | 2.31 |
| B03 | 125.68 | 3819 | 30.39 | 13.89 | 1.68 | 3.75 |
| B04 | 113.33 | 3644 | 32.15 | 12.35 | 1.48 | 2.48 |
| A06 | 108.24 | 4036 | 37.29 | 13.45 | 1.60 | 2.23 |
| B05 | 106.42 | 3340 | 31.39 | 12.04 | 1.37 | 1.77 |
| B06 | 106.01 | 3373 | 31.82 | 12.45 | 1.80 | 2.14 |
| A07 | 99.75 | 4433 | 44.44 | 12.39 | 1.51 | 1.95 |
| B07 | 66.05 | 2637 | 39.93 | 12.46 | 1.80 | 1.60 |

<sup>i</sup>Yan et al., 2021; $r^2 = 0.7$ is the default value to determine LD block by the BLINK.R package (Huang et al., 2018)

### Estimated linkage distribution (LD) and LD-decay on the Napier grass genome

A total of 65,982 genome-wide silicoDArT markers, filtered based on missing values (< 10 %), MAF (> 5 %) and known genomic position, were used to estimate linkage disequilibrium (LD) and LD-decay across the Napier grass genome. LD was analysed between pairs of SilicoDArT markers from the same AC as described previously (Muktar et al., 2019). The magnitude of *r*^2^ (square of the correlation coefficient between two markers) decreased rapidly with an increase in physical genomic distance between markers and reached a value of 0.2 at 3.48 kbp (Figure 3), which is similar to the previous estimation (2.54 kbp) that was based on the pearl millet genome (Muktar et al., 2019). The LD-decay distance varied across ACs and the slowest LD-decay was observed for A01 (7.58 kbp), while the LD decayed rapidly in B05 (1.77 kbp) and B07 (1.60 kbp). LD and LD-decay distance, and the number and density of markers used in the analysis for each AC is shown in Table 2. The proportion of pairwise *r^2^* values ≥ 0.2 (markers in LD) and ≥ 0.7 (markers in strong LD) per AC ranged from 12.04% in B05 to 15.98% in A01 and 1.37% in B05 to 2.58% in A01, respectively. The LD *r*^2^ ≥ 0.7 was the default value used to determine LD blocks by the model implemented in the BLINK.R package in GWAS analysis (Huang et al., 2018).

**Figure 3.**
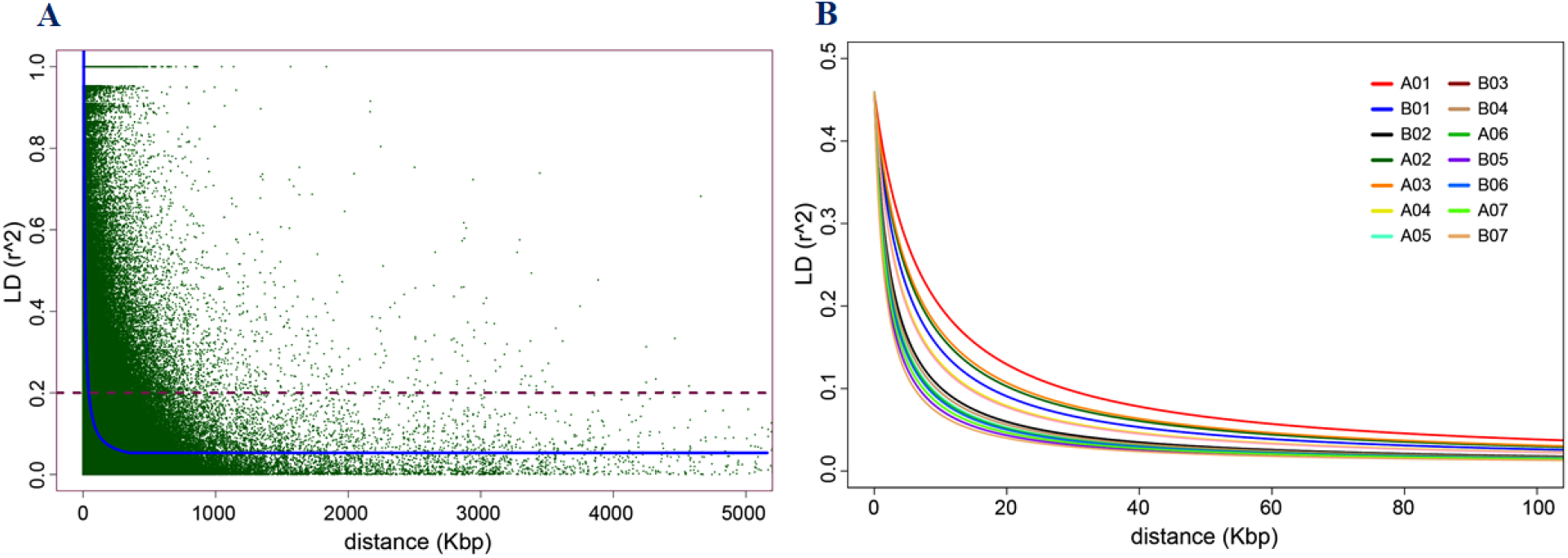
Genome-wide linkage disequilibrium (LD) decay plots. In (A), the average genome-wide LD-decay against the genomic distance (kbp). In (B), the LD-decay per assembled chromosome (AC).

### Markers associated with quantitative trait loci (QTLs) governing agro-morphological traits

A genome-wide association study (GWAS) was employed using two different mixed linear models implemented in the R packages BLINK (Huang et al., 2018) and GWASpoly (Rosyara et al., 2016). The spatially corrected phenotype values of the agronomic and morphological traits measured in the 83 Napier grass genotypes and the highly polymorphic genome-wide DArTseq markers were used. At a threshold level of *P* < 1.00E-05 in the BLINK and *P* < 0.01 (corrected for multiple testing using FDR) in the mixed linear model (MLM, using GWASpoly), more than 58 markers associated with the agronomic and morphological traits under the three-soil moisture conditions (WS-RF, DS-MWS and DS-SWS) were identified. Markers associated per trait represent different QTLs as the markers are not in LD and therefore are independent of each other.

#### WS-RF

a total of 28 markers associated with TFW, PH, LL, LW, TN, ST and IL were identified under the WS-RF condition (Figure 4A; Table 3). The 28 markers were distributed across most of the ACs, except for A02, A03 and A06. Four markers on B02, A05 and B06 were associated with TFW, two SNP markers at the bottom of B02 showed the strongest association (*P* < 1.25E-12). For PH, four markers on B01, A04 and B07 showed an association and the marker on B01 had the strongest association (*P* < 8.29E-20). Similarly, two markers on B03 and A07 and another four markers on B03, B04 and B06 were associated with LL and LW, respectively. However, the strongest association was detected on B03 for LL (*P* < 5.36E-14, by a marker IG100007430) and on B04 for LW *P* < 2.45E-20, by a marker IG100287218|F|0-5:G>C-5:G>C). The marker IG100007430 was also strongly associated (*P* < 5.38E-30) with ST. Another three markers on A01, B02 and A07 were associated with ST. For TN, one marker on A01 showed a strong association (*P* < 9.35E-18). One marker on B05 showed a strong association with IL.

**Figure 4.**
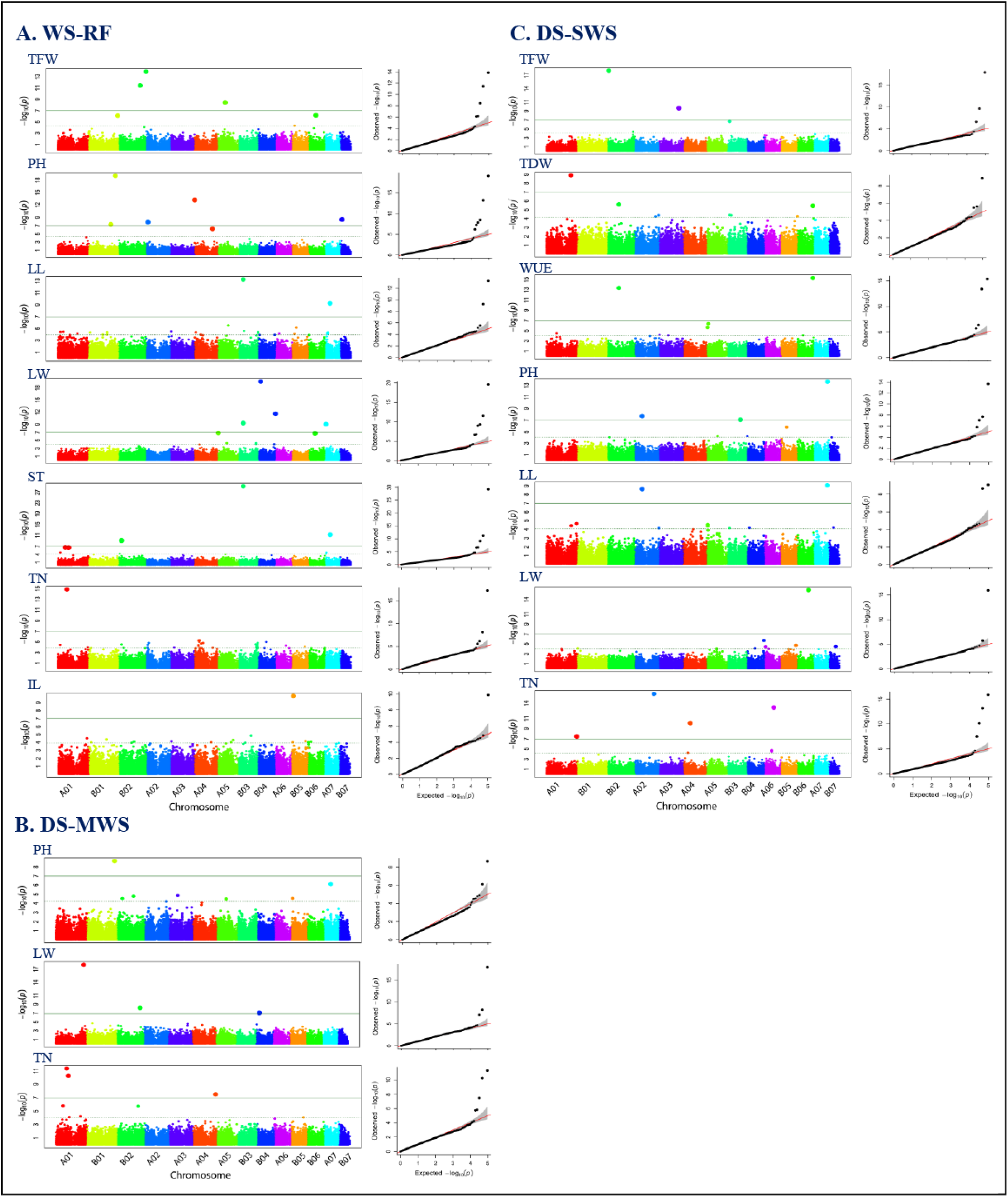
Manhattan and quantile-quantile (Q-Q) plots showing the association of markers with the agronomic and morphological traits in wet season under rainfed (WS-RF) (A), dry season under moderate water stress (DS-MWS) (B), and dry season under severe water stress (DS-SWS) (C) conditions. Markers are on the x-axis per the 14-assembled chromosome (AC). The −log10 of *P* values are plotted on the y-axis. The traits are shown at the left, total fresh weight (TFW), total dry weight (TDW), water-use efficiency (WUE), plant height (PH), leaf length (LL), leaf width (LW), stem thickness (ST), internode length (IL), and tiller number (TN).

**Table 3.** Significantly associated markers, their genomic positions, contrasting alleles, and minor allele frequency. Markers crossed the threshold levels of both the BLINK (*P* < 1.00E-05) and mixed linear model MLM (*P* < 0.01) are reported.

| Soil moisture condition | Trait | Marker | AC | Pos | allele | MAF | BLINK<br>*P*-value | MLM<br>*P*-value |
| --- | --- | --- | --- | --- | --- | --- | --- | --- |
| WS-RF | TFW | IG100206033\|F\|0-67:A>G-67:A>G | B02 | 138389691 | A/G | 0.46 | 3.65E-12 | 0.00011 |
|  |  | IG8170757\|F\|0-17:C>G-17:C>G | B02 | 176171258 | C/G | 0.30 | 1.25E-14 | 0.00013 |
|  |  | IG100035697 | A05 | 98061537 | 0/1 | 0.49 | 1.08E-09 | 7.41E-06 |
|  |  | IG100292170\|F\|0-44:C>T-44:C>T | B06 | 99625821 | C/T | 0.12 | 6.67E-07 | 0.00016 |
|  | PH | D30269880 | B01 | 143658560 | 0/1 | 0.11 | 5.23E-08 | 0.00475 |
|  |  | IG100028603 | B01 | 173322345 | 0/1 | 0.43 | 8.29E-20 | 7.52E-05 |
|  |  | IG100235976 | A04 | 675563 | 0/1 | 0.28 | 6.43E-14 | 0.00264 |
|  |  | D23616394 | B07 | 10351221 | 0/1 | 0.12 | 3.33E-09 | 0.00977 |
|  | LL | IG100007430 | B03 | 27169784 | 0/1 | 0.10 | 5.36E-14 | 4.42E-08 |
|  |  | IG100324154\|F\|0-15:C>T-15:C>T | A07 | 31359525 | C/T | 0.35 | 5.83E-10 | 0.00025 |
|  | LW | IG100007430 | B03 | 27169784 | 0/1 | 0.10 | 5.17E-10 | 3.34E-06 |
|  |  | IG100287218\|F\|0-5:G>C-5:G>C | B04 | 15150366 | G/C | 0.43 | 2.45E-20 | 0.00019 |
|  |  | IG100279889 | B04 | 112065038 | 0/1 | 0.36 | 2.64E-12 | 0.0088 |
|  |  | IG100270314 | B06 | 40015947 | 0/1 | 0.22 | 1.92E-07 | 0.0035 |
|  | TN | IG100292280\|F\|0-64:T>G-64:T>G | A01 | 58504312 | T/G | 0.06 | 9.35E-18 | 2.72E-05 |
|  | IL | IG100339862 | B05 | 9736642 | 0/1 | 0.24 | 1.32E-10 | 7.08E-05 |
|  | ST | IG100364702 | A01 | 47925751 | 0/1 | 0.36 | 2.61E-07 | 0.00100 |
|  |  | IG100222553\|F\|0-48:C>T-48:C>T | B02 | 17106838 | C/T | 0.18 | 7.66E-10 | 0.00300 |
|  |  | IG100007430 | B03 | 27169784 | 0/1 | 0.10 | 5.38E-30 | 5.47E-08 |
|  |  | IG100033657 | A07 | 32463945 | 0/1 | 0.47 | 5.49E-12 | 0.00300 |
| DS-MWS | PH | D23576564 | B01 | 179072922 | 0/1 | 0.11 | 2.22E-09 | 2.52E-05 |
|  |  | IG100244029 | A07 | 42941985 | 0/1 | 0.24 | 7.80E-07 | 2.08E-06 |
|  | LW | IG100337563 | A01 | 179359778 | 0/1 | 0.35 | 9.52E-19 | 0.00011 |
|  |  | D30269906 | B04 | 20263461 | 0/1 | 0.18 | 8.64E-08 | 0.00071 |
|  | TN | IG100224631 | A01 | 43011334 | 0/1 | 0.08 | 1.49E-06 | 1.14E-05 |
|  |  | IG100008828 | A01 | 67451249 | 0/1 | 0.18 | 4.37E-12 | 3.83E-07 |
|  |  | IG100252208 | A01 | 78125692 | 0/1 | 0.07 | 5.15E-11 | 1.16E-05 |
|  |  | D23602332\|F\|0-27:A>G-27:A>G | A04 | 144711215 | A/G | 0.26 | 3.17E-08 | 0.00023 |
| DS-SWS | TFW | D23569652 | B02 | 7364310 | 0/1 | 0.07 | 9.46E-19 | 1.66E-06 |
|  |  | IG100126885 | A03 | 125806364 | 0/1 | 0.13 | 2.28E-10 | 0.00300 |
|  |  | IG100331826 | B03 | 10779294 | 0/1 | 0.30 | 2.19E-07 | 0.00010 |
|  | TDW | D23592459 | A01 | 157193247 | 0/1 | 0.10 | 1.25E-09 | 6.08E-06 |
|  |  | IG100288904\|F\|0-22:C>G-22:C>G | B02 | 72599462 | C/G | 0.20 | 2.37E-06 | 0.00167 |

| Soil moisture condition | Trait | Marker | AC | Pos | allele | MAF | BLINK<br>P-value | MLM<br>P-value |
| --- | --- | --- | --- | --- | --- | --- | --- | --- |
|  | WUE | IG100292170\|F\|0-44:C>T-44:C>T | B06 | 99625821 | C/T | 0.12 | 3.46E-06 | 8.59E-06 |
|  |  | IG100328744 | A01 | 63184475 | 0/1 | 0.16 | 3.01E-05 | 0.00831 |
|  |  | IG100288904\|F\|0-22:C>G-22:C>G | B02 | 72599462 | C/G | 0.20 | 4.15E-14 | 0.00157 |
|  |  | IG100117610 | A05 | 6007270 | 0/1 | 0.28 | 2.07E-06 | 0.00091 |
|  |  | D23623406\|F\|0-59:G>A-59:G>A | A05 | 10548210 | G/A | 0.16 | 4.17E-07 | 0.00019 |
|  |  | IG100292170\|F\|0-44:C>T-44:C>T | B06 | 99625821 | C/T | 0.12 | 4.44E-16 | 8.22E-06 |
|  | PH | IG100218301 | A02 | 46585403 | 0/1 | 0.22 | 2.02E-08 | 7.55E-06 |
|  |  | IG100109748\|F\|0-53:C>T-53:C>T | B03 | 81852583 | C/T | 0.13 | 8.36E-08 | 0.00123 |
|  |  | D23600991\|F\|0-6:G>A-6:G>A | B05 | 34869947 | G/A | 0.28 | 1.51E-06 | 6.53E-06 |
|  |  | IG18095269\|F\|0-20:C>T-20:C>T | A07 | 89214569 | C/T | 0.28 | 2.11E-14 | 6.55E-07 |
|  | LL | IG100218301 | A02 | 46585403 | 0/1 | 0.22 | 2.31E-09 | 1.92E-08 |
|  |  | IG100009581 | A07 | 87933246 | 0/1 | 0.33 | 8.45E-10 | 7.32E-05 |
|  | LW | IG100023744 | B06 | 71606589 | 0/1 | 0.45 | 1.20E-16 | 0.00012 |
|  | TN | IG100029702 | A01 | 194732797 | 0/1 | 0.27 | 3.45E-08 | 4.55E-05 |
|  |  | IG100359692 | A02 | 123198708 | 0/1 | 0.16 | 1.32E-16 | 1.98E-05 |
|  |  | IG100219166 | A04 | 44189413 | 0/1 | 0.48 | 7.85E-11 | 0.00099 |
|  |  | IG100278405 | A06 | 59780114 | 0/1 | 0.13 | 6.75E-14 | 0.00271 |

AC = assembled chromosome; Pos. = position in a chromosome; WS-RF = wet season under rainfed condition; DS-MWS = dry season under moderate water stress condition; DS-SWS = dry season under severe water stress condition

#### DS-MWS

eight markers showed an association under moderate water stress conditions in the dry season (Figure 4B; Table 3). The eight markers were associated with PH, LW and TN and were distributed across A01, B01, A04, B04 and A07. Three markers on A01 showed the strongest association with LW (*P* < 9.52E-19) and TN (*P* < 5.15E-11).

#### DS-SWS

under severe water stress conditions in the dry season, a total of 22 markers showed an association with TFW, TDW, WUE, PH, LL, LW and TN. The markers were distributed across ACs, except for B01, B04 and B07. Two markers on B02 and B06 showed an association with the two most important agronomic traits, TDW and WUE. The strongest association with WUE was observed for two markers on B02 (*P* < 4.15E-14) and B06 (*P* < 4.44E-16). Another marker on B02 showed the strongest association (*P* < 9.46E-19) with TFW (Figure 4C; Table 3).

A total of eleven markers showed an association with the morphological traits (PH, LL, LW and TN), of which the strongest association was observed for two markers on A07 with PH and LL, one marker on B06 with LW and another three markers on A02, A04, A06 with TN. One marker on A02 showed an association with both PH and LL (Figure 4C; Table 3).

#### Common markers across the three soil moisture conditions

the MLM detected 17 markers that showed an association (*P* > −log10 =4) with traits across the three soil moisture conditions (WS-RF, DS-MWS, DS-SWS) (Table 4). Three markers on A01, A05 and an unknown position were associated with TN in all the three soil moisture conditions. Under WS-RF and DS-MWS conditions, eight common markers on B01(1), B02(3), B03 (1), B05(1) and A07(2) were associated with PH, LL, LW, TN and ST. Five common markers on B01, B04 and an unknown position showed an association with LL and TN between the WS-RF and DS-SWS conditions. Only one common marker, which was on A02 and associated with PH, was detected between the DS-MWS and DS-SWS conditions.

**Table 4.**
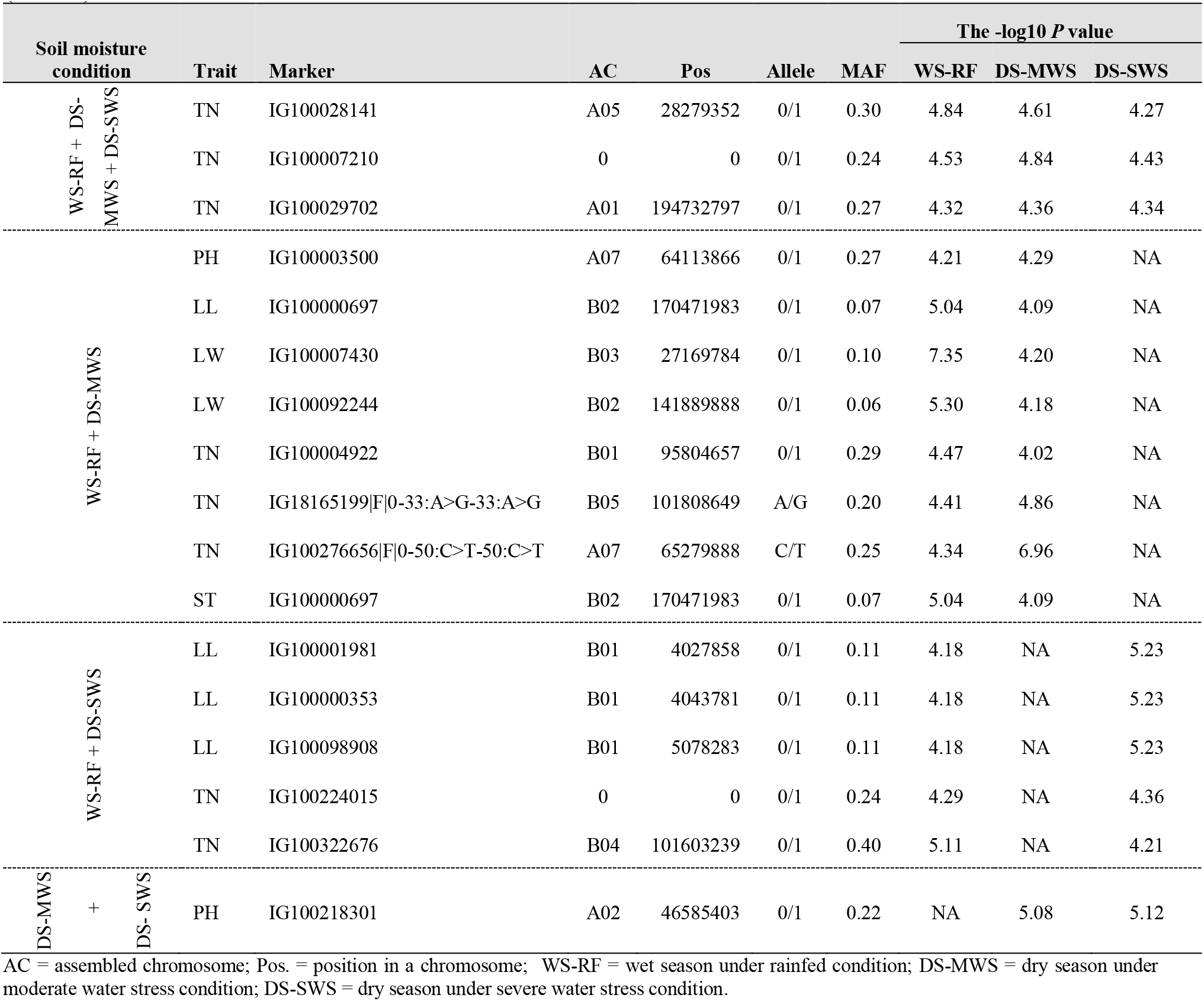
Common markers across the three soil moisture conditions. Genomic positions, contrasting alleles, minor allele frequency (MAF), and *P* values from the mixed linear model (MLM) are shown.

### Markers associated with quantitative trait loci (QTLs) governing purple pigmentation

A total of 125 markers (23 SNPs + 102 SilicoDArTs) showed an association (Figure 5A; Table 5; Supplementary Table 1) with the purple-colour pigmentation (Supplementary Figure 3) by a marker-trait association analysis using the non-parametric univariate Fisher's exact test (Warner, 2013). Of the 125 markers, 112 markers (97 SilicoDArTs and 15 SNPs) perfectly discriminated/diagnosed the seven purple genotypes. Of these markers, eight were on B01 and 19 on A03 (Table 5), while 70 were mapped only on contigs (Supplementary Table 1). However, there was no genomic position information for the remaining 15 markers (Supplementary Table 1).

**Figure 5.**
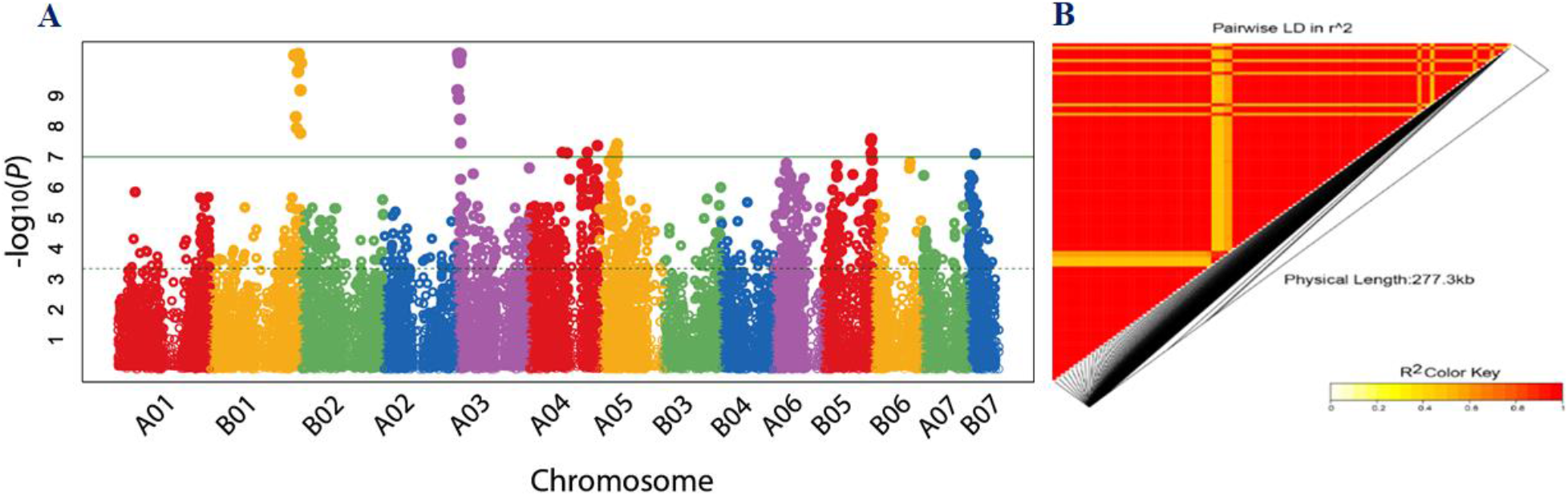
Manhattan plot showing the association of markers with purple colour (A). Markers are on the x-axis for each of the 14-assembled chromosomes (AC). The −log10 of *P* values are plotted on the y-axis. The threshold level was set at *P-*value > the −log10 value of 7. In (B), the LD heatmap for the pairwise LD of the markers associated with purple colour. As indicated by the colour key, decreasing and increasing LD is indicated by yellow and red colours, respectively.

**Table 5.** Diagnostic markers for purple colour. Markers genomic position, *P-*value, marker status, and protein associated with the marker sequences are shown. Additional diagnostic markers, but without genomic position information, are found in Supplementary Table 1.

| Marker | AC | Pos | allele | *P*-value | Status<sup>1</sup> | Protein name |
| --- | --- | --- | --- | --- | --- | --- |
| 9994171 | B01 | 190390682 | C/A | 4.39E-11 | CA | probable serine/threonine protein kinase |
| 30282538 | B01 | 181775730 | C/A | 4.71E-11 | CA/AA | NA |
| 23641768 | B01 | 185828246 | C/G | 5.05E-11 | CG | glucan endo-1,3-beta-glucosidase 3 |
| 23640854 | B01 | 195134132 | C/G | 8.39E-11 | CG/GG | sucrose transport protein |
| 10000565 | B01 | 189376936 | G/A | 1.62E-10 | GA/AA | NA |
| 100013546 | B01 | 194658440 | 0/1 | 6.59E-10 | present | bifunctional epoxide hydrolase 2-like |
| 100254243 | B01 | 184648641 | 0/1 | 4.90E-09 | present | NA |
| 100013603 | B01 | 186485183 | 0/1 | 1.14E-08 | present | cysteine-rich receptor-like protein kinase 6 |
| 100205233 | A03 | 2091285 | 0/1 | 4.39E-11 | present | F-box/kelch-repeat protein |
| 100013579 | A03 | 3641263 | 0/1 | 4.39E-11 | present | nuclear transcription factor Y subunit A-10 |
| 100013690 | A03 | 4065302 | 0/1 | 4.39E-11 | present | glycine-rich protein DOT1-like |
| 100248653 | A03 | 4447543 | 0/1 | 4.39E-11 | present | glucan endo-1,3-beta-glucosidase 3 |
| 30291207 | A03 | 5016898 | G/A | 4.39E-11 | GA/AA | Remorin family protein |
| 100013697 | A03 | 5042774 | 0/1 | 4.39E-11 | present | E3 ubiquitin-protein ligase |
| 100013640 | A03 | 5137129 | 0/1 | 4.39E-11 | present | diphosphomevalonate decarboxylase MVD2, peroxisomal-like |
| 100013671 | A03 | 5189436 | 0/1 | 4.39E-11 | present | hypothetical protein SEVIR_3G4NA32NANAv2 |
| 9897756 | A03 | 6133035 | T/C | 4.39E-11 | TC/CC | NA |
| 23641769 | A03 | 4445594 | T/C | 4.71E-11 | TC/CC | glucan endo-1,3-beta-glucosidase 3 |
| 23636057 | A03 | 5305525 | A/C | 4.71E-11 | AC/CC | NA |
| 30279984 | A03 | 3860473 | A/G | 5.05E-11 | AG/GG | NA |
| 100238305 | A03 | 5456675 | T/C | 5.05E-11 | TC | CASP-like protein 1B1 |
| 23618539 | A03 | 2845425 | C/T | 5.42E-11 | CT/TT | NA |
| 30291523 | A03 | 4425113 | C/T | 7.78E-11 | CT/TT | Cysteine-rich receptor-like protein kinase |
| 100084872 | A03 | 2094328 | 0/1 | 6.59E-10 | present | protein NUCLEAR FUSION DEFECTIVE 4 |
| 100013725 | A03 | 5362130 | 0/1 | 6.59E-10 | present | pyridine nucleotide-disulfide oxidoreductase domain-containing protein |
| 23619969 | A03 | 2925440 | C/G | 1.23E-09 | CG/GG | NA |
| 100175270 | A03 | 5573326 | T/C | 5.87E-09 | TC/CC | NA |

<sup>1</sup>Status = homozygous/heterozygous in the case of SNPs or present/absent in the case of SilicoDARts, on purple genotypes; AC = Assembled chromosome, Pos = position on AC; NA = protein name information is not available.

All the 97 SilicoDArT markers were present in the genotypes with purple colour and absent in the green genotypes. Three out of the 15 SNP markers were in a heterozygous form (for the allele associated with purple colour) in all the seven purple genotypes and were in a homozygous form (for the alternative allele) in the green genotypes. Twelve SNP markers were also in the heterozygous form (for the allele associated with purple colour) in six of the purple genotypes, however, they were in the homozygous form in one purple genotype (CNPGL_92-133-3). The markers were in very strong LD representing large haplotype blocks on B01 and A03 (Figure 5B), which were the two ACs with high synteny potentially representing the B and A’ genomes of Napier grass (Yan et al., 2021).

### Markers associated with quantitative trait loci (QTLs) governing feed quality traits

A total of 39 associated markers, one to nine markers per trait, were detected for each of the eight feed quality traits under the three-soil moisture conditions (Figure 6, Table 6). However, no associated marker was detected for CP in any of the three-soil moisture conditions.

**Figure 6.**
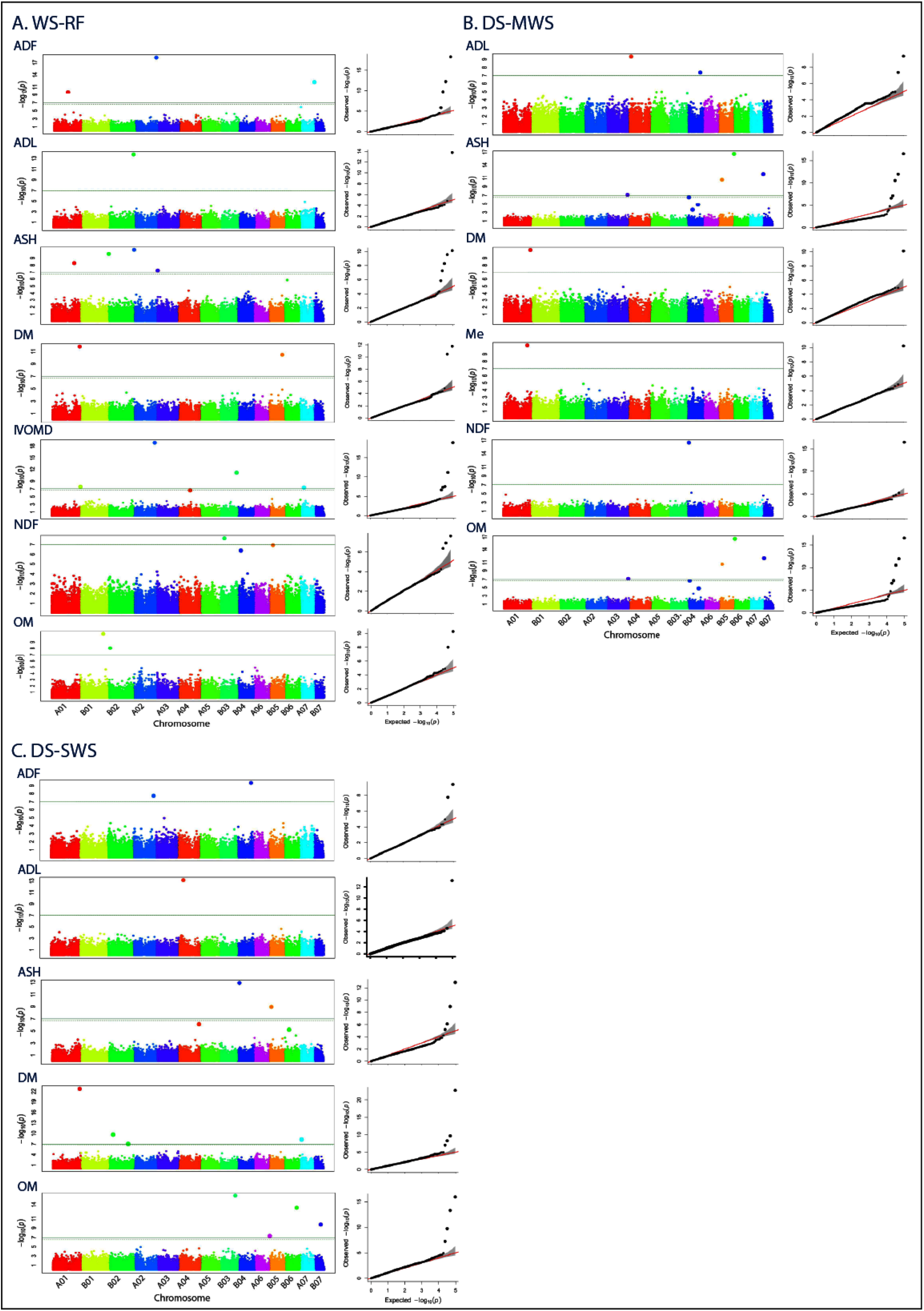
Manhattan and quantile-quantile (Q-Q) plots showing the association of markers with the nutritional quality traits in wet season under rainfed (WS-RF) (A), dry season under moderate water stress (DS-MWS) (B), and dry season under severe water stress (DS-SWS) (C) conditions. Markers are on the x-axis per the 14-assembled chromosome (AC). The −log10 of *P* values are plotted on the y-axis. The traits are shown at the left: acid detergent fibre (ADF), acid detergent lignin (ADL), ash (ASH), dry matter (DM), metabolizable energy (Me), in vitro organic matter digestibility (IVOMD), neutral detergent fibre (NDF), organic matter (OM).

**Figure 7.**
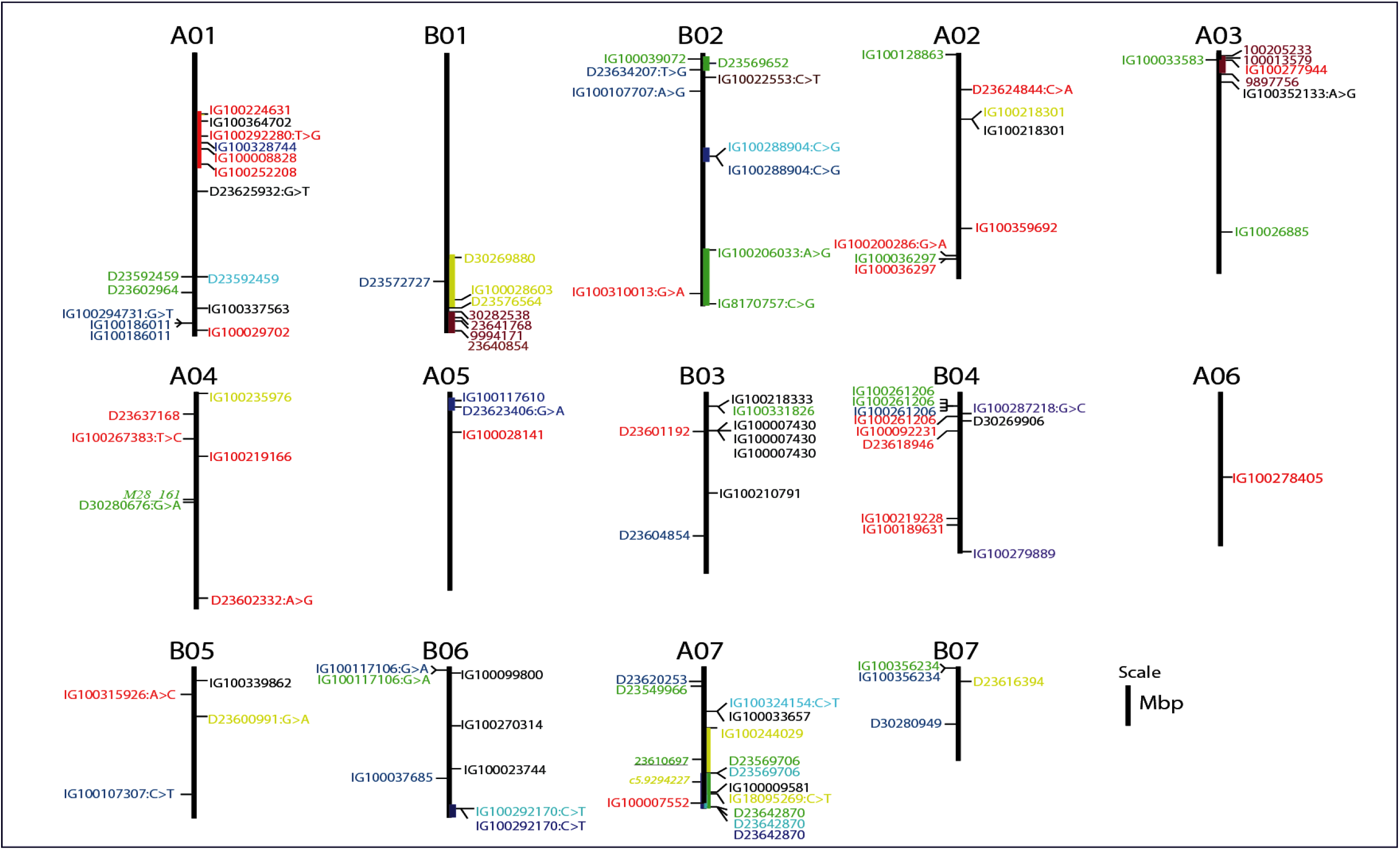
Genomic map position of markers associated with QTLs governing the agronomic, morphological, and feed quality traits. Markers associated with the agro-morphological traits are shown to the right of the assembled chromosome (AC), while markers associated with the feed quality traits are shown to the left of the AC. Markers reported previously for *Setaria italica* (Jaiswal et al., 2019; italicized) and pearl millet (Varshney et al., 2017; underlined) are also shown to the left of the AC. Markers strongly associated with TFW (total fresh weight), TDW (total dry weight), WUE (water use efficiency), PH (plant height), TN (tiller number), and PP (purple pigmentation) are shaded by green, light-blue, deep-blue, yellow, red, purple colours, respectively. Markers associated with traits that are affecting positively the feed nutritional quality (IVOMD, Me, ash) are shaded green, while markers associated with traits linked with fibre and lignin component (ADF, ADL, NDF) are shaded red. The markers associated with OM and DM are shaded blue.

**Table 6.**
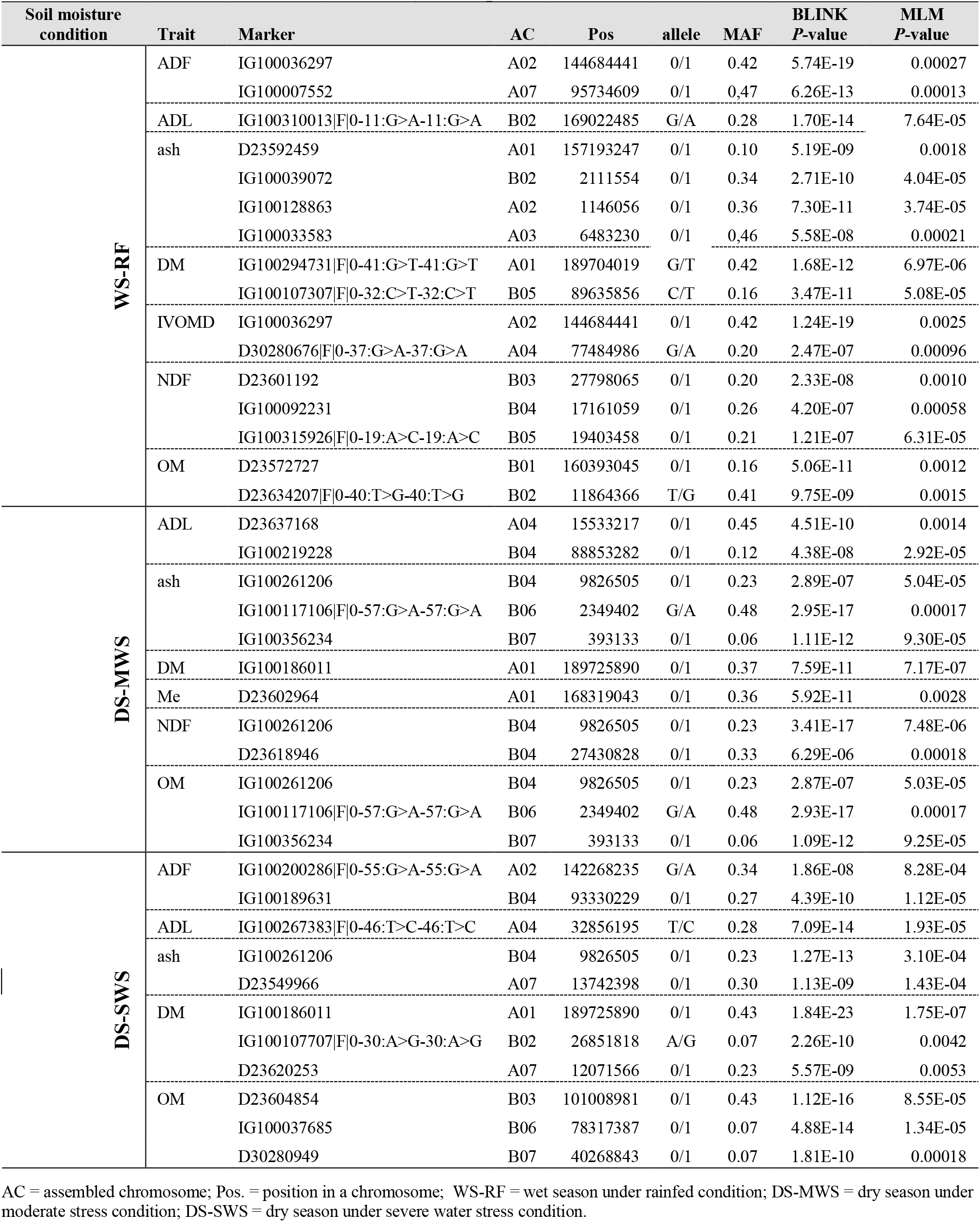
Significantly associated markers, their genomic positions, contrasting alleles, and minor allele frequency (MAF). Markers crossed the threshold levels of both the BLINK (*P* < 1.00E-05) and mixed linear model MLM (*P* < 0.01) are reported.

Under the WS-RF condition, 16 markers associated with ADF, ADL, ash, DM, IVOMD, NDF and OM were detected (Figure 6, Table 6). The markers distribution was proportional between the A’ and B genomes. Two markers on chromosome A02 and A07 were strongly associated (*P* < 6.26E-13) with ADF. The marker on chromosome A02 was also strongly associated with IVOMD. ADF and IVOMD were strongly negatively correlated (> −0.90) and hence the allele positively affecting ADF will affect IVOMD negatively and vice versa. Only one marker on chromosome B02 was associated with ADL, while four markers on chromosome A01, B02, A02 and A03 were associated with ash.

A total of twelve markers associated with ADL, ash, DM, Me, NDF and OM were detected under the DS-MWS condition (Figure 6, Table 6). No association was detected for CP, ADF and IVOMD. One marker on chromosome B04 was associated with ash, NDF and OM, in which ash was strongly negatively correlated with NDF and OM. Similarly, two markers on chromosome B06 and B07 were associated with both ash and OM. Nine out of the twelve markers were from the B genome.

Under the DS-SWS condition, eleven markers were associated with ADF, ADL, ash, DM and OM, but no association was detected for CP, Me, NDF and IVOMD (Figure 6, Table 6). Six out of the eleven markers were from the B genome.

Two markers were strongly associated with DM and ash on chromosomes A01 and B04, respectively, under both MWS and SWS conditions using both the BLINK and MLM models.

### Co-localization and functional description of sequences associated with the significant markers

Overlapping genomic positions of associated markers and co-localization with known QTLs can be used as an additional criterion to determine genomic regions controlling the traits and for the identification of candidate genes. Out of the seven markers associated with TFW, three markers that showed strong associations were on B02. Two of these markers were detected under WS-RF and were located at the bottom of B02, while the third marker, located at the top of B02, was detected in the DS-SWS condition. These marker/trait associations indicated that genomic positions from 138 mbp to 176 mbp at the bottom of B02 and at around 7 mbp at the top of B02 might harbour QTLs controlling TFW under wet and dry conditions, respectively. The two markers at the bottom of B02 were associated with genes encoding a scavenger receptor class F member 2-like and a putative box C/D snoRNA protein. The one at the top of B02 was associated with a gene encoding GDSL esterase/lipase protein (Supplementary Table 2).

The regions on B02 also contained markers associated with TDW and WUE, which were highly correlated with TFW. Two markers on B02 and B06 showed an association with both TDW and WUE. However, both these traits were strongly associated with markers at the bottom of B06 and a second stronger association at the top of A05 (Figure 6). The MLM better detected strong associations with the three highly correlated traits (TFW, TDW and WUE) at the top of B02, middle-to-bottom of A05 and the bottom of A07 (Figure 6; Supplementary Figure 8). A high level of synteny between B02 and A07, and, between A05 and B06 has been reported (Yan et al., 2021). Two markers at the bottom of A07 were associated with TFW and TDW and one of them was also associated with WUE. A pearl millet marker associated with total fresh weight (Varshney et al., 2017) was mapped at the bottom of A07 (Figure 6). The marker on B02 was associated with a gene encoding a 60 kDa jasmonate-induced protein which is involved in abiotic stress responses and defence against pathogens, while the marker on A05 was associated with a gene encoding a prolyl 4-hydroxylase protein (Supplementary Table 2).

Out of the 14 markers associated with TN under the three soil moisture conditions, five were on A01. Of these, four were close to each other and were found within the genomic region between 43 mbp to 78 mbp (Figure 6), suggesting the position of a major QTL controlling TN. Two out of the four markers were associated with genes encoding a xylanase inhibitor protein and protein DMP10 (Supplementary Table 2).

The analysis of marker-trait associations for the morphological traits was more complex compared to the agronomic traits. For PH, a greater number and stronger association were detected on B01 and A07. Genomic regions spanning approximately 29 mbp at the bottom of B01 and 46 mbp on A07 could be more important for the control of this trait. Four of the markers were associated with genes encoding a 23 kDa jasmonate-induced protein, protein SHORT-ROOT 1, F-box/FBD/LRR-repeat protein and an antifreeze protein (Supplementary Table 2). The marker on A07 was co-localized with a marker previously shown to be associated with PH in *Setaria italica* (Jaiswal et al., 2019). This region also harbours two markers associated with LL. These two markers were associated with genes encoding an ABC transporter B family member and a disease resistance protein kinase. However, the strongest association with LL was with a marker at the top of B03, which was also associated with LW and ST (Figure 6).

Genomic-position information was found for only 27 of the markers associated with the purple genotypes and all of them were mapped at the bottom of B01 and the top of A03, which are ACs with high synteny (Yan et al., 2021). This indicates that genomic regions spanning about 13 mbp at the bottom of B01 and about 5 mbp at the top of A03 harbour QTLs which govern the purple pigmentation in Napier grass. The two markers on B01 were co-localized with the markers associated with PH and another five markers on A03 were co-localized with TN and LL (Figure 6). Thirty of the markers were associated with genes encoding functional proteins (Table 5; Supplementary Table 1). One gene was annotated as an anthocyanin regulatory R-S protein and the annotation of most other genes was related to plant disease resistance.

A marker associated with ash at the top of B02 and another marker associated with ADL at the bottom of B02 were co-localized with QTL regions controlling TFW under dry and wet conditions, respectively. Another marker associated with ash at the top of A03 was co-localized with the QTL region governing the purple pigmentation (Figure 6). The genomic region spanning about 22 kbp at the bottom of A01 harbours QTLs controlling DM in all the three soil-moisture conditions. One of the markers was associated with a gene encoding a MADS-box transcription factor, which is a member of regulatory networks and governs diverse developmental processes in plants, including root development (Liu et al., 2020). A marker associated with IVOMD on A04 was co-localized with M28_161, which is a marker associated with high values of biomass digestibility in a previous study (Rocha et al., 2019).

## 4 Discussion

Genome-wide association study (GWAS) scans genetic variation across the whole genome to find signals of associations with a variation in phenotypic expression for various complex traits and is an efficient approach as it does not need the development of specific crosses, which is time consuming (Huang and Han, 2014; Flint-Garcia et al., 2003). Instead, it uses existing collections of genotypes that enables targeting a broader and more relevant genetic spectrum for plant breeders (Huang and Han, 2014). Hence, by linking markers to traits and traits to genotypes, GWAS is well suited for the genetic characterization and exploitation of genebank collections. However, for successful application of GWAS, an accurate phenotyping of the population and, as much as possible, estimating the accurate genetic value of individual genotypes is mandatory. In addition, high-density genome-wide markers and an appropriate model selection to avoid false positives are also key requirements.

### Prior and post phenotype data corrections for spatial variations

Obtaining accurate estimates of the genetic value of an individual genotype is central to the identification of markers associated with QTLs. Within an experimental field, many factors combine to generate microenvironments that vary from plot to plot and across blocks (Rodríguez-Álvarez et al., 2018), affecting biomass yield and other morphological and feed quality traits. Hence, it is important to correct for those factors when estimating genotypic effects. In this study, *a priori* and *a posteriori* spatial variability control measures were employed. As an *a priori* control measure, we used a subset of a diverse set of Napier grass genotypes, a p-rep experimental design (Williams et al., 2010) in four replications, randomization in each block, as well as check genotypes that were duplicated per block. In addition, border row plants surrounding each block were used to reduce the border effects and trait measurements were averaged from six harvests over a two-year period. However, it has been reported that these conventional control measures are not enough to account for the fine-grained spatial variability within blocks, in particular, the conventional control measures do not account for dependency between neighbouring blocks and plots within blocks, which can affect the estimation of genetic values (Elias et al., 2018; Rodríguez-Álvarez et al., 2018; Velazco et al., 2017; Lado et al., 2013). Hence, as a post data correction measure, we used the open-source model of Rodríguez-Álvarez et al. (2018) that uses two-dimensional smooth surfaces along the rows and columns of the experimental blocks to capture spatial variations. The model also allowed us to correct the phenotypes for heterogeneity in soil moisture content and soil nutrient parameters, which were used as fixed covariates in the spatial analysis. The model was able to show variation in phenotypic data due to spatial variation within a block as well as across blocks and accordingly allowed us to adjust the phenotype data, which improved the quality of our data and increased the precision in the estimation of trait heritability. A substantial improvement in heritability estimation was observed for IL, IVOMD, CP, and Me, with up to 114 %, 109 %, 90 % and 68 % improvement recorded, respectively. Similar trends of an increase in heritability estimate of traits through the application of a spatial analysis have been reported in wheat (Lado et al., 2013), sorghum (Velazco et al., 2017) and cassava (Elias et al., 2018). A greater heritability implies that a greater proportion of the phenotypic variance in the experiment was due to genetic differences among genotypes (Visscher et al., 2008), which is important for QTL identification by GWAS and other genetic analysis approaches.

### High-density genome-wide markers mapped onto the Napier grass genome

Another important prerequisite of GWAS is the availability of high-density genome-wide distributed markers, which has been a challenge especially in orphan crops which include many tropical forages. The genotyping by sequencing (GBS) method (Kilian, 2012; Jaccoud et al., 2001) applied in this study provided high density genome-wide dominant (silicoDArT) and co-dominant (SNP) markers and was found to be reliable, efficient and cost effective. The detail of the marker’s polymorphism and other related information was reported previously (Muktar et al., 2019). However, until recently we have been using the pearl millet (*Cenchrus americanus*) genome(Varshney et al., 2017) to identify the genomic positions and genome-wide distribution of the markers. Subsequently, only 17 % of the SilicoDArT markers and 33 to 39 % of the SNP markers were able to be mapped on the seven chromosomes of pearl millet (Muktar et al., 2019; Paudel et al., 2018). Fortunately, a Napier grass reference genome was made public recently (Yan et al., 2021) which enabled us to generate much more information, including genomic position and genome-wide distribution for most of the markers and an enhanced estimation of LD and LD-decay in the Napier grass genome. More than 90 % of the SNP and 73 % of the SilicoDArT markers were able to be mapped onto the fourteen assembled chromosomes (AC) of the Napier grass genome. The markers were evenly distributed across the genome with some gaps around the middle part of each AC, which probably represents the gene-poor regions of centromeres (Varshney et al., 2017; Bennetzen et al., 2012). However, the density and distribution of the markers was by no means comprehensive and this resource does not represent the extensive genome coverage required to detect all the possible QTLs in the Napier grass genome. Rather it represents about a quarter of the estimated marker-density and QTL detection power in Napier grass, as estimated using the average LD-decay across the genome previously (Muktar et al., 2019) and in the present study. In this study, similar to the previous report (Muktar et al., 2019), the LD decayed very rapidly, on average at about 3.48kbp, and varied across the ACs. A long LD block was observed in A01, while the shortest one was in B07. Interestingly, the number of markers in each AC was proportional or positively correlated with the average LD-decay distance per AC. Detailed information about LD and LD-decay across the Napier grass genome and LD variation between collections can be found in Muktar et al. (2019).

### Association analysis and correction for population structure and cryptic relatedness

The third important concern in GWAS is the presence of population structure and cryptic relatedness in the mapping panel that could prevent the association analysis from correctly identifying the true marker-trait association and could lead to the identification of false-positive or spurious associations (Huang and Han, 2014; Flint-Garcia et al., 2003). Therefore, it is important to include population structure and pairwise kinship matrix as covariates and to select the appropriate statistical models that can sufficiently deal with those factors. The detail of population structure and genetic diversity in our Napier grass collections was reported previously (Muktar et al., 2019), in which five to seven subpopulations were detected using different genetic diversity study approaches. A similar population stratification was seen in the 84 genotypes used in this study (Supplementary Figure 9).

For the morphological, agronomic, and feed quality quantitative traits, we employed two different GWAS models implemented in the recently reported Bayesian-information and Linkage-disequilibrium Iteratively Nested Keyway (BLINK) model (Huang et al., 2018) and the mixed linear model (MLM) that takes into account both population structure and pairwise kinship matrix to avoid false positives (Rosyara, et al., 2016). The BLINK model is based on linkage disequilibrium (LD) and controls confounding issues arising due to cryptic relatedness and multiple testing corrections by using all tested markers within an LD block (Huang et al., 2018). Furthermore, the model takes population stratification information as covariates and hence the first three to five PCs from a PCA analysis were used in our analysis. The model efficiency in controlling the above-described confounding factors was demonstrated by the quantile-quantile (Q-Q) plots, which showed a similar distribution of observed and expected *p*-values along the diagonal line for most of the markers and a sharp curve at the end of the line representing a few associated markers. This model identified more than 58 and 39 independent markers associated with the agro-morphological and nutritional quality traits, respectively, under the three different soil moisture conditions. The MLM identified a much greater number of associated markers, however, most of the associated markers per trait were in LD with each other and with the most associated marker and represent a single or few QTLs aligned with the trait.

The two models were complementary to each other in that both identified many common markers and a few different markers associated with the traits. The BLINK model identified a single most associated marker per QTL that helped us to estimate the genomic position of QTLs for a trait, while many associated markers per QTL region were detected by the MLM and that was important for further analysis and identification of more candidate genes and their annotation within the QTL region. The MLM also identified common markers associated with a trait across the three soil moisture conditions. There were a few QTL regions detected only by the BLINK but not by the MLM, and vice versa, which could be attributed to the differences in algorithms and parameters used in the two models (Huang et al., 2018; Rosyara, et al., 2016). However, we employed both models, together with the different algorithms and parameters, to optimize the selection of QTL regions and candidate genes for further exploration.

### Markers associated with forage biomass and water-use efficiency traits

GWAS identified six QTL regions associated with TFW, which represents the above-ground fresh biomass production. The QTLs were at the top and bottom of B02, the bottom of A03, and the top of A05, B03 and B06. Particularly, the QTLs on B02 could be more interesting for breeding applications as they showed a strong association and were detected by both the BLINK and the MLM models. The QTLs at the top of B02 were associated with increased biomass yield under severe water stress conditions in the dry season, while those at the bottom were associated with high biomass production during the wet season. The QTLs at the bottom of B02 were linked with genes encoding scavenger receptor and box C/D snoRNA proteins, which are proteins involved in immune response and rRNA modification (Streit et al., 2020). In our previous marker-trait analysis (Habte et al., 2020), we reported markers associated with annual dry weight yield at the bottom of A03 (14 markers), B05 (14 markers) and B02 (4 markers).

Three QTL regions towards the bottom of A01, and, around the middle of B02 and B06, showed an association with TDW and, specifically, increased total dry matter production under the SWS conditions in the dry season. The QTLs on B02 and B06 were also associated with improved WUE, which is the most important factor in improving productivity under limited water availability in the dry season and in addressing the current challenges to forage production due to climate change. An additional QTL associated with WUE was detected at the top of A05, which shows high synteny and co-linearity with the bottom of B06 (Yan et al., 2021). Hence, we can speculate that these two ACs may represent the A’ and B homeologous chromosomes of Napier grass. Napier grass is an allotetraploid grass (2n=4x=28) with a complex genome (A’A’BB), with the A’ genome showing a high degree of homology with the pearl millet (2n = 2x = 14 with AA genomes) A genome (dos Reis et al., 2014). The QTL on B02 was linked with a gene encoding a 60 kDa jasmonate-induced protein which is involved in abiotic stress responses and defence against pathogens (Rustgi et al., 2014), while the marker on A05 was linked with a gene encoding a prolyl 4-hydroxylase protein, which is involved in plant growth and development and in responses to abiotic stresses (Vlad et al., 2007). The marker on A05 was mapped on pearl millet linkage group 2, probably co-localized with the major drought-tolerant QTL (DT-QTL) reported in pearl millet (Tharanya et al., 2018).

The MLM detected additional QTL regions at the bottom of A07, which show high synteny with the upstream part of B02 (Yan et al., 2021). The MLM also detected association peaks, in which several consecutive markers showed an association, at three genomic regions: the top of B02, bottom of A05, and, A07. These three QTL regions were strongly associated with the three most highly correlated traits (TFW, TDW and WUE) indicating the tight linkage between the agronomic and water use efficiency traits. Co-mapping and tight linkage of QTLs for agronomic and water use efficiency traits were reported in pearl millet, in which close linkages between QTLs controlling the traits in four genetic regions of linkage group 2 were detected (Tharanya et al., 2018). The possibility of a pleiotropic effect, in which the three traits are controlled by a single QTL, was reported in *Setaria italica* (Feldman et al., 2018). Co-localization of QTLs and the markers associated with them can potentially be exploited in marker-assisted selection to develop high biomass producing Napier grass varieties for dry and water deficit areas.

### Markers associated with morphological traits

In this study, a greater number of QTL regions associated with the morphological traits (PH, LL, LW, ST, IL and TN) were detected by a total of 35 markers, which might suggest that the genetic architecture is more complex when compared to the architecture of the agronomic traits. Among these, the QTL region at the top of A01 was strongly associated with TN, especially under wet and moderate water stress conditions, and hence is considered a major QTL. Higher tillering ability is an important trait in establishment and regrowth in grasses (Pereira et al., 2013) and was positively correlated with the TFW, TDW and WUE traits while showing no correlation with traits linked with fibre and lignin (ADF, NDF and ADL), indicating its importance in the high biomass production of Napier grass without compromising the feed nutritional quality.

We also identified two QTL regions for PH at the bottom of B01 and A07, one QTL region at the top of B03 controlling LL, LW and ST, and another QTL region for IL at the top of B05. The markers associated with the QTLs will be useful in a Napier grass improvement program through the application of marker-assisted selection. The QTL region for PH on B01 is co-localized with the QTL region associated with purple pigmentation. A similar result was reported in pearl millet (Azhaguvel et al., 2003), in which loci controlling purple foliage colour, dwarf plant height, and resistance to downy mildew and rust diseases were co-localized on pearl millet linkage group 4. The QTL region associated with PH on A07 was co-localized with a marker associated with PH in *Setaria italica* chromosome 5 (Jaiswal et al., 2019), which shows a high degree of synteny and co-linearity with the top part of the Napier grass A07 (Yan et al., 2021).

By using sequence tags corresponding to the associated markers, we identified several candidate genes linked to the QTL regions. Two markers in the QTL region associated with TN were linked with genes encoding a xylanase inhibitor protein and a protein DMP10. In the QTL region associated with PH, four markers were linked with genes encoding a 23 kDa jasmonate-induced protein, protein SHORT-ROOT 1, F-box/FBD/LRR-repeat protein and an antifreeze protein. These proteins are involved in abiotic stress responses and in different developmental process, for example, the SHORT-ROOT protein has been reported to play a role in regulation of primary, lateral, and adventitious root developments in Arabidopsis (An et al., 2019; Rustgi et al., 2014; Lucas, et al., 2011). The candidate genes will be useful for further characterization of the QTL regions and potentially cloning of the QTLs using the candidate gene association mapping approach.

### Markers associated with purple pigmentation

For the qualitative purple pigmentation trait, QTL regions were identified by the non-parametric univariate Fisher's exact test (Warner, 2013) that detected a greater number of significantly associated markers around two genomic regions. Two QTL regions towards the bottom of B01 and the top of A03 were associated with purple pigmentation in Napier grass. The two genomic regions show a high synteny block (Yan et al., 2021) representing the B and A’ homeologous chromosomes of Napier grass (dos Reis et al., 2014). Out of the different expanded genes involved in anthocyanin biosynthesis in Napier grass, phenylalanine ammonia lyase (PAL) and chalcone synthase (CHS) were detected on B01 (Yan et al., 2021). One of the markers associated with the QTLs was linked with a gene annotated as an anthocyanin regulatory protein and the annotation of most other genes within the QTL regions was related to plant disease resistance, suggesting the co-localization of the two traits. Anthocyanins are naturally occurring pigments belonging to the group of flavonoids, a subclass of the polyphenol family (Martin et al., 2017), and are involved in biotic and abiotic stress tolerance (Naing et al., 2018). In addition, anthocyanin expression is associated with the expression of stress response genes in plants, such as genes involved in drought, water logging, cold tolerance and disease resistance (Ren et al., 2019).

Our findings are in line with previous reports on pearl millet (Varalakshmi et al., 2012; Azhaguvel et al., 2003), in which they mapped the *P* foliage colour locus to pearl millet linkage group 4. The pearl millet linkage group 4 shows strong synteny and co-linearity with the Napier grass A03 and also some synteny with B01 (Yan et al., 2021). In addition, downy mildew and rust resistance genes have been identified in this genomic region of the pearl millet genome (Varalakshmi et al., 2012; Azhaguvel et al., 2003). According to Varalakshmi et al. (2012), the pearl millet purple pigmentation of the leaf sheath, midrib and leaf margin are co-inherited under the control of a single dominant locus (the ‘midrib complex’) and are inseparably associated with the locus governing the purple coloration of the internode.

The markers identified in this study can be used as tagging markers for map-based cloning of the QTLs controlling purple coloration (anthocyanin) and stress tolerance related genes, which will be very important for the genetic improvement of Napier grass. The markers might also be useful in marker-assisted selection for disease resistance and stress tolerance in Napier grass.

### Markers associated with feed quality traits

A total of eleven markers, representing different QTL regions across the genome, were identified for IVOMD, Me and ash under the three-soil moisture conditions. These traits were strongly correlated and positively affect the feed nutritional quality. No associated marker was identified for CP, however, this trait was strongly correlated with IVOMD and Me, and hence the markers identified for these two traits could be used in marker-assisted selection for CP. Marker M28_161 that was associated with high values of biomass digestibility in a previous study (Rocha et al., 2019) was co-localized with the marker (D30280676|F|0-37:G>A-37:G>A) associated with IVOMD on A04, in this study. Similarly, thirteen markers were detected for ADF, NDF and ADL, which are mainly linked with the cell wall components of fibre and lignin that negatively affect the feed nutritional quality. Improving the digestibility of forages by reducing lignin and fibre is a major goal in forage crop breeding programs (Habte et al., 2020; Rocha et al., 2019).

Most of the markers associated with the traits that positively affect feed nutritional quality were from the Napier grass A’ genome, while the markers associated with the traits linked with fibre and lignin components were mainly from the B genome, which might indicate a functional differentiation between the two sub-genomes as suggested previously (Yan et al., 2021; Zhang et al., 2020). However, QTL regions overlapping for the positive and negative traits were detected at the bottom of A02 and top of B04. These regions need to be further investigated to determine whether these overlapping regions were caused by pleiotropy.

## Conclusions

1. In this study, we determined the true genetic response of Napier grass genotypes to the morphological, agronomic and feed quality traits by using different approaches that reduced the environmental effects and errors as much as possible. We also showed that the quality of the phenotype data and precision in the estimation of trait heritability can be improved using spatial analysis.
2. Most of the high-density genome-wide markers generated in this study were mapped onto the recently assembled Napier grass genome. Hence, genomic position information was generated for more than 90 % of the SNP and 73 % of the SilicoDArT markers, which improved the identification of genomic regions that harbour QTLs for the important forage traits. The availability of the genome sequence is an important asset and provides a lot of opportunities for the characterization and cloning of the QTLs and for the development of improved Napier grass varieties.
3. Our study led to the identification of at least 58 novel QTL regions and associated markers for morphological, agronomic and water use efficiency traits, and more than 39 novel QTL regions for feed quality traits, and, provided clearer insights into the genetic architecture of the traits in Napier grass. The desirable alleles of the associated markers identified in this study will be useful in Napier grass improvement through marker-assisted breeding. In addition, candidate genes linked with the associated markers were identified and can be used for further characterization and validation of the QTLs through candidate gene association mapping. Further validation of the associated markers, QTLs and the candidate genes identified here may lead to a better understanding of the genetic/genomic bases for the trait’s genetic variation.
4. We also identified two major QTL regions associated with the Napier grass purple colour, which is associated with anthocyanin pigmentation. The markers identified in this study can be used as tags for map-based cloning of the QTLs controlling purple pigmentation and stress tolerance related genes, which will be very important for the genetic improvement of Napier grass. Anthocyanin expression is associated with the expression of stress-response genes, such as genes involved in drought, cold and waterlogging tolerance, and disease resistance.
5. The information gained from the present study will be useful for the genetic improvement of Napier grass production with enhanced water use efficiency while maintaining its nutritional quality.

## Acknowledgements

The research was conducted in Feed and Forage Development (FFD) program at the ILRI forage genebank in Addis Ababa and at the Bishoftu field site, Ethiopia. The authors would like to thank EMBRAPA for making their germplasm and breeding lines available for the study through the Africa Brazil Agricultural Innovation Marketplace Project. Authors would also like to thank Dr. Raphael Mrode and Dr. Jorge Fernando Pereira for reading, commenting, and correcting the manuscript.

## Author Contributions

C.S.J. designed and supervised the project and the manuscript writing, M.S.M. analyzed the data and wrote the manuscript, E.H. collected the phenotype data, A.T. and A.T.N. collected leaf samples, A.T. extracted DNA, Y.A. involved in phenotype data collection, K.W.L. involved in the supervision of the phenotyping project. J.Z. involved in the supervision of the genotyping project. All authors made a significant contribution to the development of this manuscript and approve it for publication.

